# Alternative start codon selection shapes mitochondrial function during evolution, homeostasis, and disease

**DOI:** 10.1101/2025.03.27.645657

**Authors:** Jimmy Ly, Yi Fei Tao, Matteo Di Bernardo, Ekaterina Khalizeva, Christopher J. Giuliano, Sebastian Lourido, Mark D. Fleming, Iain M. Cheeseman

## Abstract

Mitochondrial endosymbiosis was a pivotal event in eukaryotic evolution, requiring core proteins to adapt to function both within the mitochondria and in the host cell. Here, we systematically profile the localization of protein isoforms generated by alternate start codon selection during translation. We identify hundreds of pairs of differentially-localized protein isoforms, many of which affect mitochondrial targeting and are essential for mitochondrial function. The emergence of dual-localized mitochondrial protein isoforms coincides with mitochondrial acquisition during early eukaryotic evolution. We further reveal that eukaryotes use diverse mechanisms—such as leaky ribosome scanning, alternative transcription, and paralog duplication—to maintain the production of dual-localized isoforms. Finally, we identify multiple isoforms that are specifically dysregulated by rare disease patient mutations and demonstrate how these mutations can help explain unique clinical presentations. Together, our findings illuminate the evolutionary and pathological relevance of alternative translation initiation, offering new insights into the molecular underpinnings of mitochondrial biology.

## Introduction

Mitochondrial function is critical to eukaryotic cells and dysregulation of mitochondrial processes has been linked to hundreds of rare human diseases^1^ ^2^, impacting millions of individuals globally. The evolution of the mitochondria required specialized adaptations in the early eukaryotic ancestor^3^ ^4^ ^5^ ^6^. Core biological processes, including DNA replication, transcription, and translation, must occur within both mitochondria and the host cell, requiring the trafficking of the protein machinery involved in these processes to multiple, distinct cellular compartments. To ensure the proper localization of essential activities within mitochondria, most proteins are directed via a primary import pathway that relies on mitochondrial targeting signals present in the protein’s N-terminus of the protein^7^ ^8^. These signals facilitate the translocation of unfolded proteins into the mitochondria through the TOM complex^7^. The import of proteins into the mitochondria is typically irreversible such that protein molecules imported into the mitochondria cannot re-localize to other organelles. This creates a substantial challenge when a protein is also required within the cell cytoplasm or nucleus. One evolutionary strategy to achieve similar protein activities in each compartment is through a gene duplication event, followed by the acquisition of a mitochondrial targeting signal for one paralog^3^ ^5^ ^9^. For example, topoisomerase activity is crucial for preventing supercoiling during DNA replication for both mitochondrial and nuclear DNA. In humans, this function is carried out by *TOP1* and *TOP1MT,* highly related genes that share 85% sequence similarity. TOP1 localizes to the nucleus whereas TOP1MT localizes to the mitochondria through its acquired mitochondrial targeting signal^10^. Instead of a gene duplication event, the alternative decoding of a single gene by controlling transcription, splicing, or translation which has the potential to produce multiple protein isoforms, including those with differing localization^11,12^ ^13^ ^14^ ^15^ ^16^. In particular, protein isoforms with distinct N-terminal sequences would be particularly likely to affect mitochondrial targeting, protein secretion, and other subcellular trafficking events that rely on signal sequences^17^. However, our understanding of the molecular mechanisms that generate proteins capable of functioning across multiple cellular locations remains limited.

Human cells generate significant proteomic diversity through alternative start codon selection, with an estimated 50% of mRNAs producing more than one protein product^18–20^. Mechanisms such as leaky ribosome scanning^21^ ^22^ ^15^, internal ribosome entry ^23^ ^24^, initiation at near-cognate start codons^25^ ^12^ ^26^ ^27^, and uORF reinitiation^28^ ^29^, amongst others^30^ ^31^ ^32^, can each drive the use of downstream alternative start codons within a single mRNA. Downstream translation initiation that is in-frame with the annotated start codon will result in N-terminally truncated protein products. Furthermore, although the region upstream of an annotated start codon is typically described as an 5′ untranslated region (5′ UTR), these sequences can also encode functional protein products. In addition to uORFs, the presence of in-frame alternative start codons in the 5′ UTR, including non-canonical start codons such as CUG and GUG, can produce N-terminal extensions to the annotated protein. However, the widespread functional significance and roles of these alternative isoforms remain largely unexplored, with these alternative start sites often discounted as translational noise.

In this study, we reveal that alternative start codon selection serves as a pivotal mechanism regulating mitochondrial function, evolution, and dysregulation. Through a systematic analysis of alternative N-terminal variants generated in human cells, we show that these isoforms frequently exhibit distinct subcellular localization compared to their annotated counterparts, with a substantial fraction exhibiting altered mitochondrial localization. We further trace the evolutionary origins of alternative start codon selection to primitive eukaryotes following mitochondrial endosymbiosis. Finally, we identify multiple dual-localized alternative translational isoforms for mitochondrial genes that are specifically mutated in rare human diseases. These findings highlight the essential role of alternative translation initiation in understanding mutations linked to human disease and offer new insights into the evolution and molecular underpinnings of mitochondrial biology.

## Results

### Pervasive changes in the localization of alternative N-terminal isoforms

To define the contributions of alternative translational isoforms that differ from the annotated isoform by their N-terminus, we sought to explore the consequences of alternative initiation site usage. Using ribosome profiling, our recent work empirically defined translation initiation sites empirically in human HeLa cells identifying 15903 initiation sites in 7736 genes^33^. Of the identified genes, 4495 genes express a single N-terminal isoform, whereas 2697 produce two or more N-terminal isoforms including in-frame N-terminal extensions or truncations (Fig. 1A; Fig. S1A-C). As a protein’s N-terminus is often critical to direct its subcellular localization, we evaluated genes with one or more protein variant for differences in localization signal sequences including mitochondrial targeting sequences and signal peptides for protein secretion (Fig. 1B).

**Figure 1.**
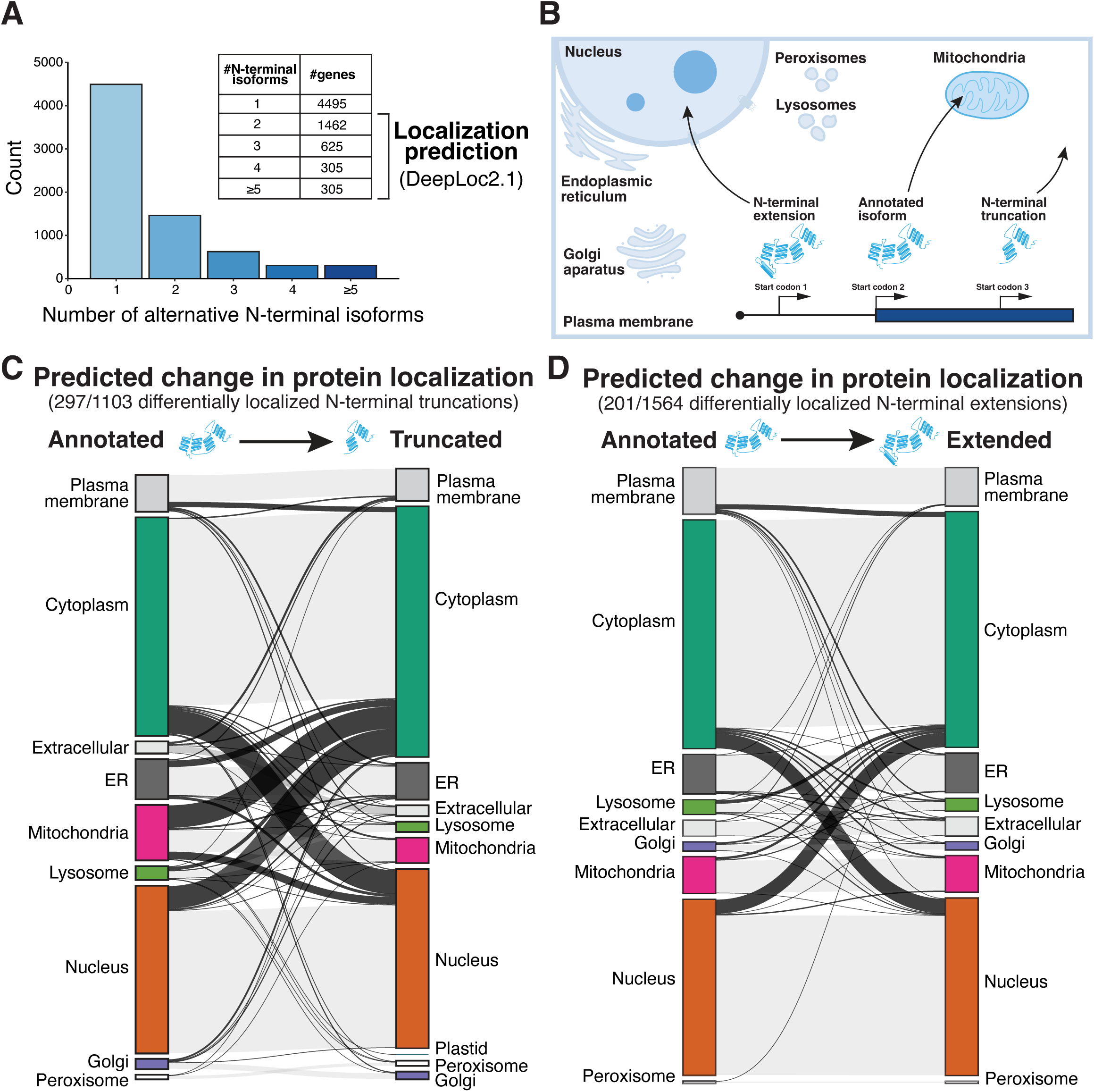
Alternative start codon selection regulates protein subcellular localization. (**A**) Histogram of the number of alternative N-terminal protein isoforms for per gene in HeLa cells. Note that some genes have multiple “annotated” N-terminal isoforms. (**B**) Schematic highlighting the importance of the proteins N-terminus in subcellular localization and how alternative start codon selection may generate differentially localized protein isoforms. (**C**) Sankey plot showing the change in predicted subcellular localizations between the annotated isoform and N-terminal truncation based on DeepLoc2.1. The connecting black lines represent predicted changes in localization between annotated and truncated protein isoforms. Faint colored lines indicate no pairs of isoforms that have the same predicted localization. The thickness of the lines represents relative frequency. (**D**) As described in Fig. 1C except depicting the localization changes between the annotated and N-terminal extension.

To systematically investigate the subcellular localization of alternative N-terminal isoforms, we used DeepLoc2.1, a subcellular localization prediction algorithm based on protein language models ^34^. Using DeepLoc2.1, we predicted the localization of genes products with both an annotated ORF and an alternative N-terminal extension or truncation (Fig. S1D and S1E). For genes with multiple annotated or alternative N-terminal extensions or truncations start codons, we selected the protein isoform from the ORF type with the highest translation initiation efficiency based on our previous quantitative ribosome profiling data (Fig. S1D). Among 1103 pairs of annotated proteins and N-terminally truncated isoforms, 297 (26.9%) were predicted to display differential localization. Similarly, of 1564 pairs of annotated proteins and N-terminal extensions, 201 (12.9%) were predicted to show distinct localization patterns (Fig. 1C-D; Supplemental table S1). Strikingly, of the differentially localized alternative translational isoforms, 108 (21.7%) represented changes in mitochondrial localization (Fig. 1C-D; Supplemental table S1), suggesting that mitochondrial genes may have particularly evolved alternative translation to generate protein products to function in multiple cellular compartments.

To validate the differential localization of alternative translational isoforms experimentally, we generated pairs of tagged constructs for 40 genes, tagging either the annotated coding sequence or the alternative isoform with a C-terminal GFP (Fig. 2A; Fig. S1F-G and S2A-F; Supplemental table S2). Of the 40 pairs of translational isoforms tested, 27 pairs showed differential subcellular localization, as described in more detail below (Fig. 2A; Fig. S1G and S2A-F; Supplemental table S2). For several selected genes, we also tested whether dual localization could be observed using expression of a construct designed to mimic the native mRNA including the endogenous 5′ UTR region. Ectopic expression of constructs containing the 5′ UTR – coding sequence (CDS) – GFP for TRNT1, MRPS38, and GARS1 each resulted in dual mitochondrial and nuclear/nucleolar localization (Fig. S2A-D). Importantly, we were able to deconvolve this dual localization through expression of a single translational isoform (Fig. 2A, 2E; Fig. S2A-D). Together, our data suggest that alternative translation initiation generates a substantial number of protein variants with distinct subcellular localizations, with a particular impact on mitochondrial localization.

**Figure 2.**
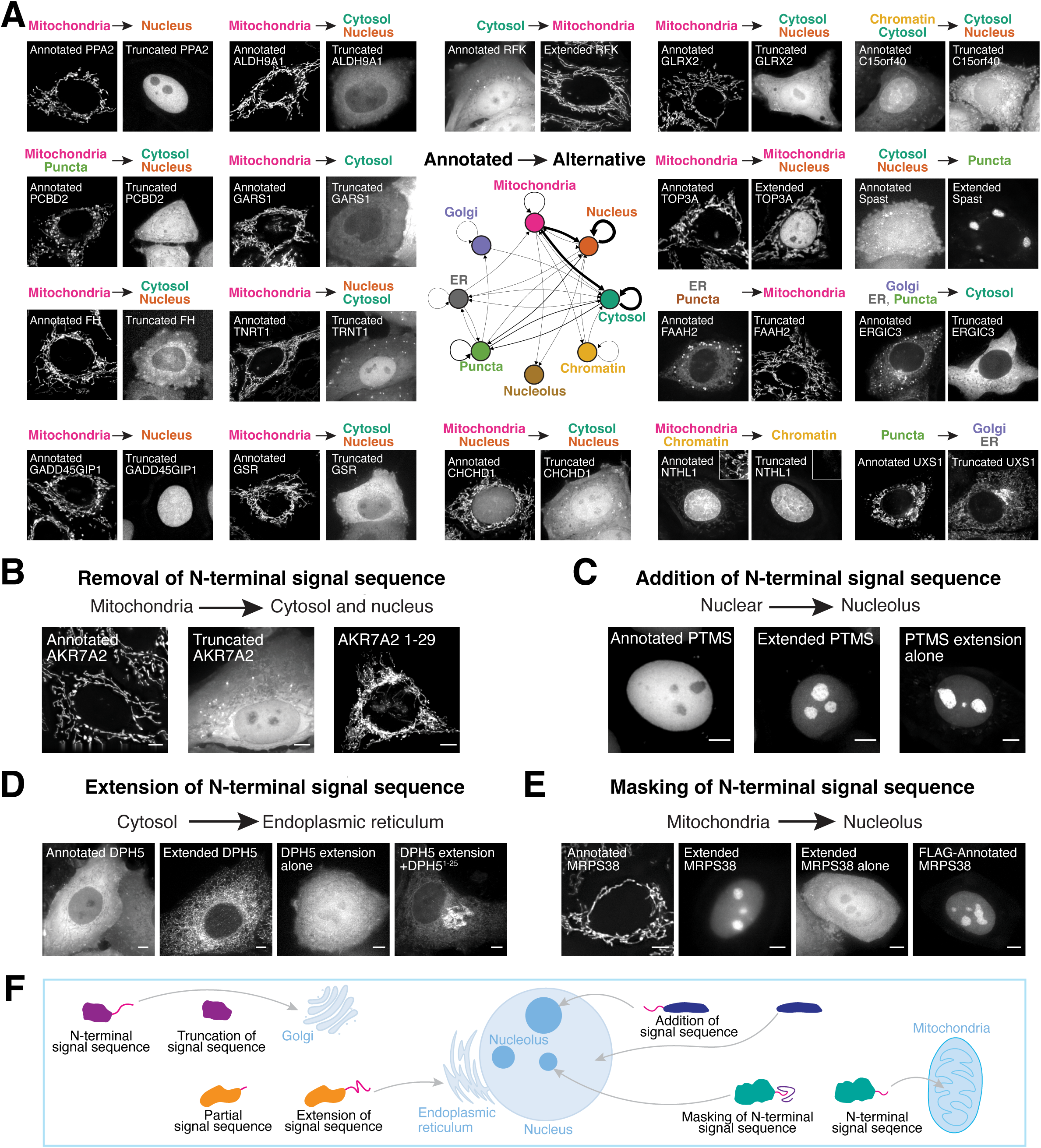
Pervasive differential localization of alternative N-terminal isoforms. (**A**) Live-cell images of indicated genes from the isoform specific localization screen. Summary of the differential localization of alternative N-terminal isoforms is shown in the localization network graph in the middle. The line thickness represents increased frequency of this change in localization between isoforms of a gene. Inset images are scaled to be brighter than full size image. (**B**) Left, live cell imaging of annotated AKR7A2-GFP shows localization to mitochondria. Middle, live cell imaging of truncated AKR7A2-GFP shows localization to cytosol and nucleus. Right, live cell imaging of AKR7A2^1–29^-GFP shows localization to mitochondria. (**C**) Left, live cell imaging of annotated PTMS-GFP shows localization to nucleus. Middle, live cell imaging of extended PTMS-GFP shows localization to nucleolus. Bottom, live cell imaging of PTMS N-terminal extension alone tagged with GFP shows localization to nucleolus. (**D**) Live cell imaging of annotated DPH5-GFP (cytosol), extended DPH5-GFP (ER), DPH5 extended region alone tagged with GFP (cytosol), and DPH5 N-terminal extension with the first 25 amino acids of annotated DPH5 tagged with GFP (ER). (**E**) Live cell imaging of annotated MRPS38/AURKAIP1-GFP, extended MRPS38/AURKAIP1-GFP, MRPS38/AURKAIP1 extended region alone tagged with GFP, N-terminally FLAG tagged annotated MRPS38 with a C-terminal GFP. Scale bar, 5 µm. (**F**) Schematic representation of the mechanisms by which alternative start codon selection may alter subcellular localization.

### Mechanisms of differential alternative N-terminal isoform localization

We next sought to define the molecular basis by which the subcellular localization of these protein variants is altered. We identified several core strategies that act to create changes in protein localization. First, we identified N-terminal truncations in which the absence of annotated N-terminal amino acids selectively eliminates a signal sequence responsible for targeting the annotated protein to a specific compartment. For example, the genes AKR7A2, TRNT1, PNPO, GSR, and CMPK1 each contain a downstream alternative translation initiation site that removes an N-terminal targeting signal, leading to altered localization (Fig. 2B; Fig. S2G). Second, we identified N-terminal extensions to the annotated protein that contain the complete sequence elements needed to create a new localization signal. For example, we identified an N-terminal extension in Riboflavin kinase (RFK) that is sufficient to encode a mitochondrial targeting signal (Fig. S2H) and an extension in Parathymosin (PTMS) containing a nucleolar localization signal, each of which result in distinct subcellular localization compared to their annotated isoforms (Fig. 2C; Fig. S2I). Third, we identified N-terminal extensions that complement incomplete signal sequences at the N-terminus of the annotated protein. For example, the N-terminally extended isoform of DPH5 localizes to the endoplasmic reticulum, whereas the annotated isoform resides in the cytosol (Fig. 2D). Neither the N-terminal extension alone nor the first 25 amino acids of the annotated DPH5 sequence is sufficient to form a complete signal peptide (Fig. 2D; Fig. S2J). However, when these two regions are combined, they generate a functional signal sequence that results in endoplasmic reticulum localization (Fig. 2D; Fig. S2J). Fourth, we identified N-terminal extensions that alter protein subcellular localization by “masking” or “internalizing” the annotated N-terminal targeting signal, thereby preventing it from being recognized. For example, the annotated isoform of AURKAIP1/MRPS38 localizes to mitochondria via its N-terminal targeting signal (Fig. 2E; Fig. S2K). However, the N-terminally extended MRPS38 isoform localizes to the nucleus/nucleolus, likely because the extension displaces the mitochondrial targeting signal away from the N-terminus, disrupting its function (Fig. 2E). Indeed, we found that adding an unrelated N-terminal sequence (FLAG-S tag) to the annotated MRPS38 isoform redirected it to the nucleus/nucleolus (Fig. 2E). Similarly, the N-terminally extended TOP3A isoform localizes to the nucleus instead of mitochondria through the internalization of its mitochondrial targeting sequences (Fig. S2L). This work highlights the diversity of mechanisms by which alternate translational isoforms alter protein localization to allow a single gene to generate multiple protein isoforms with distinct subcellular localizations (Fig. 2F).

### Early eukaryotic origins of alternative translation initiation and dual localization

Amongst the alternative translational isoforms that displayed dual localization, the largest fraction involved changes in mitochondrial protein localization (Fig. 1C-D and Fig. 2A). The emergence of mitochondria was a pivotal event in eukaryotic evolution. In the early eukaryotic ancestor, DNA, protein translation, protein folding, and antioxidant activities needed to be independently maintained in both the mitochondria and other compartments within the cell ^3^ ^4^ ^5^ ^6^. Our results suggest that one way in which cells enable these critical dual-localized protein functions in both the mitochondria and other compartments is through the generation of alternate protein isoforms. If this is the case, we reasoned that the existence of these alternate isoforms would be broadly conserved across eukaryotes, arising after mitochondrial acquisition and the streamlining of the mitochondrial genome (Fig. 3A).

**Figure 3.**
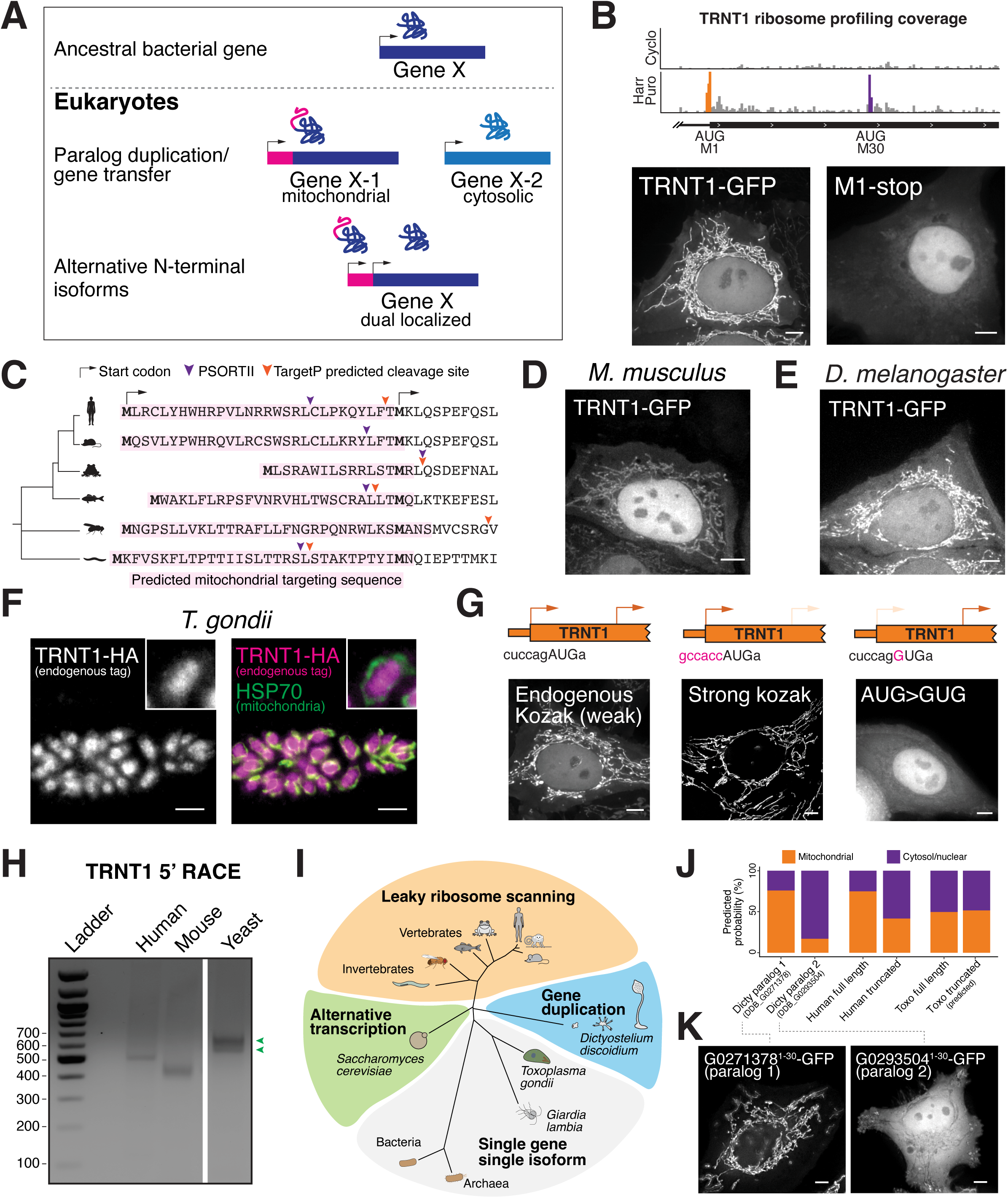
Early eukaryotic evolution of alternative start codon selection. (**A**) Schematic of how alternative start codon selection to generate dual-localized protein isoforms may replace paralog duplication/gene transfer during mitochondrial acquisition. (**B**) Top, ribosome profiling reads around the TRNT1 start codons from HeLa cells. Orange bars represent ribosome protected fragments at the annotated TRNT1 start codon whereas purple reads represent reads at the alternative start codon. Bottom, live cell imaging of cells expressing wild type TRNT1 5′ UTR – CDS – GFP (left) or TRNT1 5′ UTR – CDS – GFP but with a premature stop codon between Met1 and Met30. (**C**) TRNT1 protein sequence alignment around the truncated start codon in Metazoans. The sequence with magenta shade represents the predicted mitochondrial targeting signal from DeepLoc. The arrowheads indicate the predicted mitochondrial processing peptidase cleavage site by the indicated program. (**D**) Live cell imaging of *M. Musculus* TRNT1 5′ UTR – CDS – GFP in HeLa cells expressed from the safe harbor locus. (**E**) Live cell imaging of *D. Melanogaster* TRNT1 5′ UTR – CDS – GFP in HeLa cells expressed from the safe harbor locus. (**F**) Immunofluorescence imaging of endogenously tagged TRNT1 – mCherry – HA in *T. Gondii*, co-stained with the HSP70 mitochondrial marker. Inset region indicated by white box. (**G**) Live cell imaging of human TRNT1 with the indicated start codon mutants. (**H**) 5′ rapid amplification of cDNA ends of TRNT1 from several organisms. Green arrow head indicates different 5′ mRNA products for TRNT1 in *S. Cerevisiae*. White gap indicates image crop. (**I**) Cladogram showing the presence of TRNT1 alternative start codon selection and the underlying mechanism. (**J**) DeepLoc2 localization predictions for the two TRNT1 paralogs in *Dictyostelium discoideum*, the alternative TRNT1 isoforms from human, and *T. gondii* TRNT1 from the first and next downstream methionine. (**K**) Live cell imaging of *D. discoideum* G0271378 and G0293504 N-terminal region tagged with C-terminal GFP, expressed in HeLa cells. Scale bar, 5 µm.

By analyzing evolutionary conservation (PhyloP) ^35^ across vertebrates, we found that the start codons for N-terminal truncations of mitochondrial genes are more highly conserved compared to the start codons N-terminal truncations from non-mitochondrial genes (Fig. S3A-C), suggesting an evolutionary pressure to retain these dual-localized mitochondrial genes. For example, we identified conserved alternative translation initiation sites in glycine-tRNA synthetase (GARS1) in human HeLa cells, mouse embryonic stem cells ^18^, and budding yeast ^36^ that produce dual mitochondrial and cytosolic isoforms (Fig. S3D-E). The mechanism that drives the production of alternative GARS1 isoforms is evolutionarily conserved from humans to budding yeast ^36^, occurring through leaky ribosome scanning due to a weak start codon at Met1 initiation site and downstream translation initiation (Fig. S3D-E). Similarly, we observed a strong conservation of alternative isoforms for TRNT1, which facilitates the maturation of both mitochondrial and nuclear-encoded tRNAs by adding a CCA sequence to their 3’ ends ^37^ (Fig. S3F-G). To perform this dual function in distinct cellular compartments, we found that human TRNT1 undergoes alternative translation initiation at two sites: Initiation at Met1 (AUG) that produces a mitochondrial-localized isoform, and initiation at Met30 (AUG) that generates a nuclear/cytosolic isoform (Fig. 3B). This mechanism is widely conserved across species (Fig. 3C), including mice (Fig. 3D), flies (Fig. 3E), and budding yeast ^38^. These results suggest that there is a strong conservation of these alternative translational isoform across eukaryotes.

Alternatively, we reasoned that organisms that have dispensed aspects of mitochondrial function would not have a selective pressure to retain this dual localization behavior. For example, *Trypanosoma brucei* ^39^ and the apicomplexan parasite *Toxoplasma gondii* ^40^ have functional mitochondria but lack mitochondrial-encoded tRNAs and instead import processed nuclear-encoded tRNAs into the mitochondria. In addition, the primitive eukaryote *Giardia lambia* lacks mitochondria altogether ^41^. Consistent with the absence of a requirement for mitochondrially-localized TRNT1 in these species, we found that the alternative TRNT1 initiation site is absent in *G. lambia*, *T. brucei*, and *T. gondii* (Fig. S3H), suggesting the production of a single nuclear-localized TRNT1 translational isoform. To test this model directly, we monitored the subcellular localization of *T. gondii* TRNT1 (TGGT1_315810). We found that TRNT1 endogenously tagged with a C-terminal mCherry-HA localizes to the nucleus but not to mitochondria in *T. gondii* (Fig. 3F), aligning with prior *T. gondii* mitochondrial proteomics data ^42^. These observations suggest that the alternative translation initiation sites in TRNT1 evolved from its ancestral bacterial/archaeal enzyme in early eukaryotes, likely after mitochondrial acquisition but only in organisms that have retained mitochondrially-encoded tRNAs. These results highlight the importance of alternative translation in generating dual-localized protein variants for mitochondrial function and eukaryotic evolution (Fig. 3A).

### Diverse evolutionary pathways for production of dual-functional proteins

Our results indicate that a subset of dual-localized mitochondrial/cytosolic isoforms are deeply conserved, coinciding with mitochondrial endosymbiosis. We next sought to evaluate the gene regulatory strategies that control dual protein production across species. One major pathway of alternative start codon selection is through leaky ribosome scanning that allows for the usage of multiple start sites in a single mRNA ^32^. For example, in vertebrates, TRNT1 is transcribed as primarily a single mRNA transcript (Fig. S3I-J). TRNT1 alternative isoform translation occurs via leaky ribosome scanning due to a weak Kozak context at Met1 (Fig. 3G; Fig. S3G) such that strengthening Met1 context favors the mitochondrial isoform, whereas mutating Met1 (AUG>GUG) produces only the cytosolic/nuclear isoform (Fig. 3G).

The decoding a single gene into multiple protein products can also through alternative transcription initiation. In this case, the use of distinct promoter sites can result in mRNAs of differing lengths such that the shorter product lacks the ability to use the upstream translation start site. Indeed, in budding yeast (*S. cerevisiae* ^38^) the TRNT1 homolog CCA1 produces two distinctly localized isoforms through alternative transcription initiation rather than leaky scanning (Fig. 3H-I; Fig. S3K). Lastly, instead of encoding a dual-localized TRNT1 as a single gene, the slime mold *Dictyostelium discoideum* has two TRNT1 paralogs with distinct predicted localizations. One paralog (DDB_G0271378) contains a strong N-terminal MTS and is predicted to localize to the mitochondria, whereas the other (DDB_G0293504) lacks an N-terminal MTS and is predicted to localize to the nucleus (Fig. 3I-J; Fig. S3L-N). Indeed, based on expression in human cells, we found that the N-terminus of DDB_G0271378 functions as a mitochondrial targeting signal whereas the DDB_G0293504 paralog did function as a signal sequence (Fig. 3K). These findings highlight the diverse gene regulatory strategies that act across evolution to maintain the production of dual-localized activities for a single enzyme to create both mitochondrial and nuclear populations (Fig. 3I).

### 5′ UTR length regulates dual-localized N-terminal isoform selection

Although established mechanisms that influence start codon selection, such as leaky scanning, non-AUG translation, RNA structure, uORF reinitiation, alternative transcription initiation, and alternative splicing can explain the generation of most alternative protein isoforms in our analysis, we identified several alternative start codons within single mRNA isoforms whose usage could not be clearly explained by known translational control pathways. For example, AKR7A2 produces differentially localized isoforms: translation from the first AUG results in mitochondrial localization, whereas initiation at a downstream AUG led to cytosolic/nuclear localization (Fig. 2B). However, a AKR7A2 5′ UTR – CDS – GFP construct expressed under a dox-inducible promoter resulted in only mitochondrial localization, suggesting the exclusive usage of the first AUG (Fig. 4A). In contrast, we observed dual localization for AKR7A2 following the transfection of a similar *in vitro* transcribed AKR7A2 5′ UTR – CDS – GFP mRNA, consistent with the behavior of endogenous AKR7A2 (Fig. 4A).

**Figure 4.**
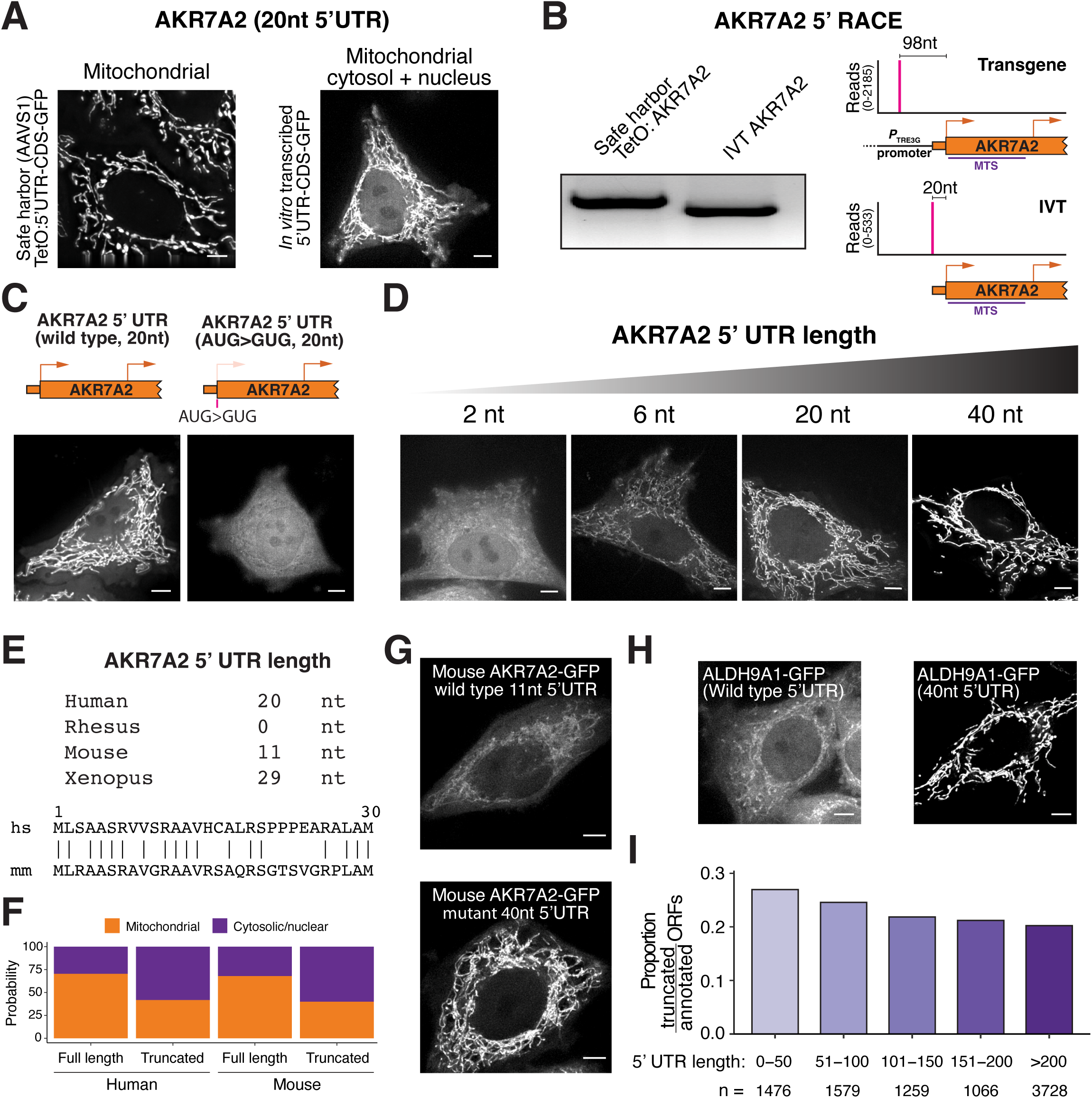
5′ UTR length regulates alternative isoform selection. (**A**) Live cell imaging of AKR7A2 5′ UTR – CDS – GFP in HeLa cells using two different systems. Left, transgene was inserted into the AAVS1 locus under a dox inducible promoter. Right, reporter was *in vitro* transcribed, capped, and polyadenylated then transfected into HeLa cells. (**B**) Left, 5′ RACE of endogenous AKR7A2 from transgenic AKR7A2 expressed under dox inducible promoter and *in vitro* transcribed AKR7A2 mRNA. Right, quantification of 5′ RACE reads by sequencing. (**C**) Left, Live cell imaging of AKR7A2 after transfecting mRNA containing the AKR7A2 5′ UTR – CDS – GFP. Right, localization of AKR7A2 where the annotated start codon was mutated to GUG. (**D**) Live cell imaging of AKR7A2 5′ UTR – CDS – GFP with various 5′ UTR lengths. (**E**) Top, AKR7A2 5′ UTR length humans, monkey, mouse, and frog. Bottom, protein sequence alignment for AKR7A2 from human and mouse. (**F**) DeepLoc2 predictions for full length and truncated AKR7A2 from human and mouse. (**G**) Live cell imaging of HeLa cells transfected with *in vitro* transcribed mouse AKR7A2 5′ UTR – CDS – GFP reporters with different 5′ UTR lengths. (**H**) Live cell imaging of ALDH9A1 with *in vitro* transcribed wild type (left) and long (right) 5′ UTR – CDS – GFP mRNA. Scale bar, 5 µm. (**I**) Bar plot showing the relative proportion of N-terminal truncations to main ORFs for genes in different 5′ UTR bins.

To understand the difference in transgene localization, we considered whether this was due to the addition of nucleotides in the 5′ UTR (which resulted from expression from the dox-inducible promoter (*P*_TRE3G_ ^43^) (Fig. 4B). To investigate the role of 5′ UTR length, we transfected human cells with *in vitro*-transcribed mRNAs encoding the native AKR7A2 5′ UTR (20 nt), a chimeric AKR7A2 construct containing the *Xenopus* beta-globin 5′ UTR (40 nt), and truncated AKR7A2 5′ UTRs (6 nt and 2 nt), each coupled to the full AKR7A2 coding sequence and a C-terminal GFP tag. The native 20-nt AKR7A2 5′ UTR produced both mitochondrial and cytosolic/nuclear protein isoforms (Fig. 4C). Mutating the first start codon (AUG>GUG) completely eliminates production of mitochondrial localized protein (Fig. 4C), such that the ratio of mitochondrial vs. nuclear/cytosolic isoform represents the relative production from the two start codons. In contrast to the wild type AKR7A2 20-nt 5′ UTR, the extended 40-nt 5′ UTR produced isoforms localized exclusively to mitochondria, suggesting the usage of the first but not second start codon (Fig. 4D). Reciprocally, further shortening the 5′ UTR led to a shift favoring cytosolic/nuclear localization over mitochondrial localization (Fig. 4D). Importantly, we found that this short 5′ UTR length and production of dual-localized AKR7A2 is conserved across mammals (Fig. 4E and F). Indeed, transfection of the mouse AKR7A2 mRNA yielded similar results as human AKR7A2 mRNA, suggesting evolutionary pressure to maintain its short 5′ UTR for dual decoding (Fig. 4G). We observed a similar 5′ UTR length-dependent behavior for the dual decoding of ALDH9A1 alternative N-terminal isoforms, which has an endogenous 5′ UTR length of only 4 nucleotides (Fig. 4H; Fig. S4A-B). These results highlight the critical role of short 5′ UTRs in alternative start codon selection and dual isoform localization.

To test whether short 5′ UTR lengths are correlated with the production of alternative translational isoforms, we analyzed the relationship between 5’ UTR length and downstream translation initiation. We found that mRNAs with short 5’ UTRs (≤50 nt) exhibited more downstream translation initiation resulting in N-terminal truncations compared to those with longer 5’ UTRs (>50 nt), suggesting a broad paradigm (Fig. 4I; Fig. S4C-D, Supplemental table S3). Amongst other biological processes, we additionally found that mitochondrial mRNAs are enriched for short 5′UTRs (Fig. S4E-H). Thus, we propose that 5′ UTR length regulation could contribute to diverse biological processes by expanding the functional human proteome, with a particular role in enabling mitochondrial function through the production of dual-localized isoforms. This model is consistent with prior structural models and synthetic reporter mRNAs that propose increased leaky ribosome scanning on unusually short 5′ UTRs^44^ ^45^ (Fig. S4I).

### Alternative translational isoforms contribute to dual protein activities

The widespread conservation of these alternative protein isoforms highlights their potential functional significance, but such dual-localized isoforms (and their corresponding functions) would be overlooked if only the annotated isoform were considered. To evaluate these functional contributions, we first tested AURKAIP1/MRPS38, which contains a 28-amino-acid N-terminal extension that initiates from a CUG start codon (Fig. S5A-D). The MRPS38 extension results in nucleolar localization through masking the annotated N-terminal mitochondrial targeting signal (Fig. 2E). MRPS38 has an established role as a core subunit of the small mitochondrial ribosome ^46^, but a nucleolar role has not been suggested in prior work. We confirmed the alternate usage of this upstream CUG start codon using a combination of mass spectrometry (Fig. S5B), immunofluorescence (Fig. S5C), and luciferase reporter assays (Fig. S5D). To assess the functional contribution of the extended AURKAIP1/MRPS38 isoform to cellular fitness, we generated isoform-specific AURKAIP1/MRPS38 indel alleles using CRISPR/Cas9-based gene editing and performed competitive growth assays (Fig. 5A). As expected, the annotated isoform of MRPS38 is required for cell growth due to its essential role as a component of the mitochondrial ribosome (Fig. 5B). In addition, selectively eliminating the nucleolar CUG-initiated AURKAIP1/MRPS38 isoform resulted in a significant growth defect, suggesting that the extended isoform also plays a critical role in cell viability (Fig. 5B).

**Figure 5.**
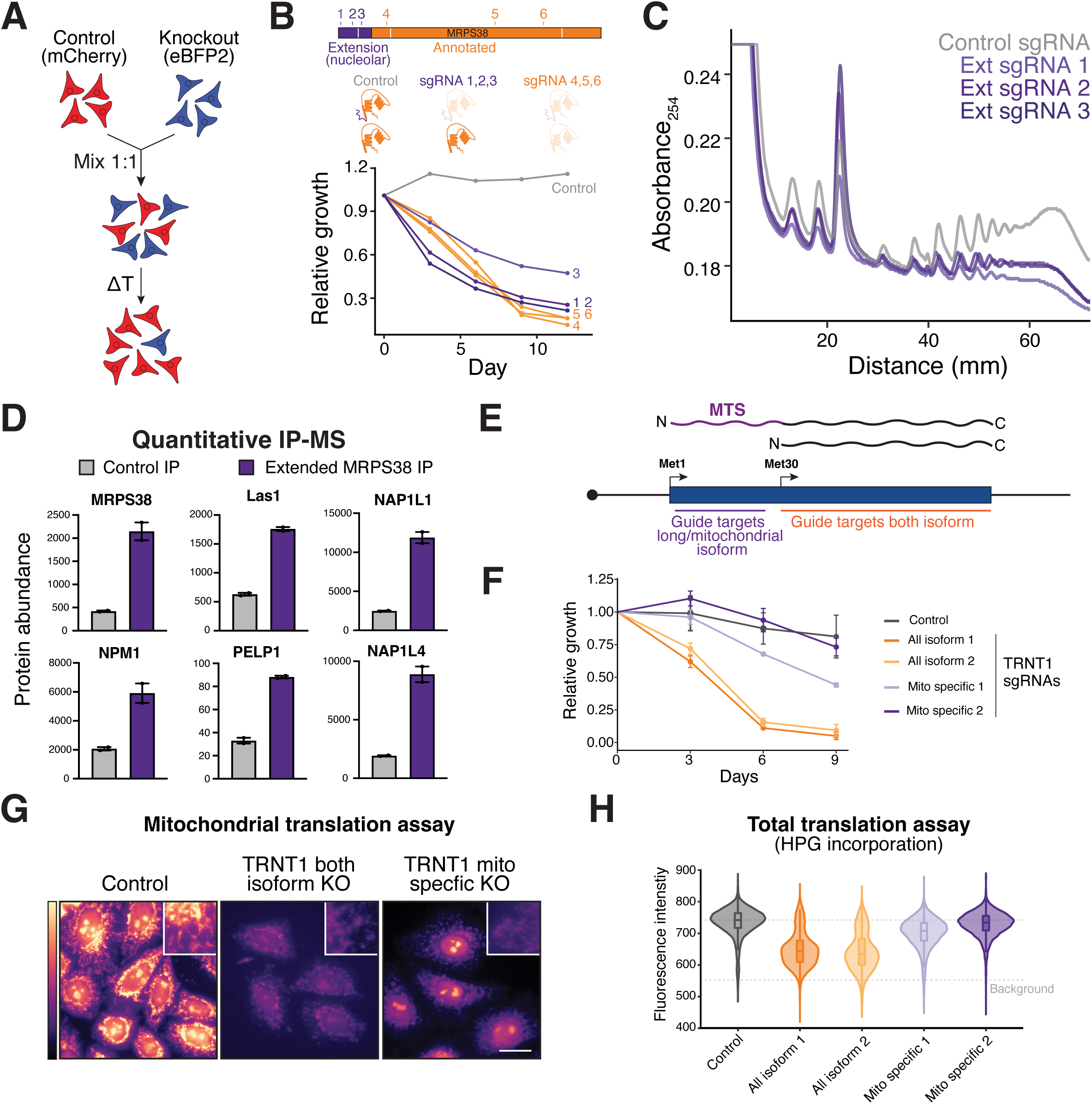
Alternative start codon selection promotes dual protein function. (**A**) Schematic of competitive growth assay. (**B**) Competitive growth assays in cells expressing various gRNAs to selectively deplete MRPS38 isoforms (top). Guide RNAs (1-3) specifically eliminating the CUG-initiation MRPS38 extension result in a growth defect (bottom). (**C**) Sucrose gradient profiles from day 6 control and MRPS38 extension knockout cells. (**D**) Quantitative MS analysis of extended MRPS38/AURKAIP1 immunoprecipitation reveals interactions with several nucleolar and nuclear proteins. (**E**) Schematic model for designing isoform specific CRISPR guide RNAs for genes with N-terminal truncations. (**F**) Competitive growth assays in cells expressing various gRNAs to selectively deplete TRNT1 isoforms. (**G**) Imaging of mitochondrial translation rate assay in control and isoform specific TRNT1 knockouts. Inset region, representing a zoom in on mitochondrial stain is indicated by white box. Scale bar, 25 µm. (**H**) Violin plot showing flow cytometry results from total HPG incorporation assay in control and isoform specific TRNT1 knockouts. The dotted line represents median HPG intensity for control knockout cells (top dashed line) and cells without HPG (bottom dashed line, background defined by a no HPG control).

We hypothesized that the extended AURKAIP1/MRPS38 isoform may be important for cytosolic ribosome activity given its localization to the nucleolus. To test this, we used CRISPR-Cas9 to specifically eliminate the extended AURKAIP1/MRPS38 isoform and analyzed cytosolic translation using polysome gradient analysis. Consistent with a role in promoting cytosolic ribosome activity, these isoform-specific knockouts exhibited a reduced cytoplasmic polysome/monosome ratio (Fig. 5C) suggesting that the extended isoform contributes to cytosolic ribosome biogenesis within the nucleolus. In addition to a functional role, we found that the extended AURKAIP1/MRPS38 isoform interacted with several nucleolar proteins, including LAS1-like ribosome biogenesis factor (LAS1), Nucleophosmin 1 (NPM1), and Proline-, Glutamic acid-, and Leucine-rich protein 1 (PELP1), as well as other nuclear proteins such as Nucleosome assembly protein 1-like 1 (NAP1L1) and 4 (NAP1L4) (Fig. 5D; Fig. S5E, Supplemental table S4). This is consistent with a role for nucleolar MRPS38 in ribosome biogenesis.

We next examined molecular basis for the usage of the essential upstream alternative CUG start codon in MRPS38. Analysis of the AURKAIP1/MRPS38 5′ UTR sequence revealed a strong RNA secondary structure downstream of the CUG codon ^47^, which is predicted to pause the scanning ribosome and promote CUG initiation (Fig. S5F, ^12^ ^48^ ^49^ ^50^). Disrupting this structure with silent mutations reduced CUG-based translation in reporter assays (Fig. S5G) and only produced a mitochondria-localized protein isoform (Fig. S5H), confirming the loss of the CUG-initiated nucleolar isoform. These findings suggest that the AURKAIP1/MRPS38 mRNA is decoded as two distinct isoforms: one localized to the mitochondria, where it supports mitochondrial translation, and another localized to the nucleolus, where it is required for efficient cytosolic translation. Thus, our findings demonstrate that alternative translation creates differentially localized protein isoforms that account for the essential biological functions attributed to individual genes.

Beyond AURKAIP1/MRPS38, we identified multiple other genes that are predicted to rely on alternative N-terminal isoforms to fulfill their established functions. For instance, TRNT1 has documented roles both within and outside the mitochondria ^37^. However, the annotated isoform is exclusively mitochondrial, suggesting that the proposed nuclear/cytosolic functions may instead be mediated the N-terminal truncated isoform (Fig. 2). To test this, we used CRISPR-Cas9 cutting between the two translation initiation sites to selectively eliminate the upstream translational isoform while preserving initiation of the downstream isoform (Fig. 5E). As a comparison, we used Cas9 to target a region downstream of both start codons to abolish both protein variants. We found that guide RNAs targeting both TRNT1 isoforms conferred greater fitness defects than those disrupting only a single isoform (Fig. 5F). In addition, we monitored both mitochondrial ^51^ and total translation using L-Homopropargylglycine (HPG) incorporation ^52^. We found that guide RNAs eliminating both TRNT1 isoforms reduced both global translation and mitochondrial translation, whereas guides targeting the longer TRNT1 isoform only eliminated mitochondrial translation (Fig. 5G-H). Overall, these results highlight the importance of alternative start codon selection in generating multiple protein isoforms each of which make important contributions to cellular function and viability.

### Multiple differentially localized alternative translational isoforms are mutated in rare disease

Our analysis highlights the essential role of alternative protein isoforms that enable dual activities both within and outside the mitochondria. Given the importance mitochondrial function for human health, we next assessed whether there are patient mutations that affect specific N-terminal isoforms in rare genetic diseases (Fig. 6A). To identify pathogenic mutations that target a specific alternative translational isoform for a gene, we analyzed the ClinVar database ^53^. Of the 3,334,918 ClinVar variants examined, we found 24,183 isoform-specific mutations associated with rare diseases. This includes 22,487 missense, 660 nonsense, and 1,036 frameshift mutations (Fig. 6B; Fig. S6A), with 258 (24%) of genes containing a nonsense or frameshift mutation that is predicted to eliminate the annotated but not truncated isoform of a gene (Supplemental Table S5). Of the 22 pairs of annotated and N-terminal truncated isoforms analyzed here (Fig. 2; Supplemental table S2), 9 genes (FH, PPA2, GARS1, GSR, LAGE3, NAXE, NTHL1, PNPO, TRNT1) had a nonsense or frameshift mutation between the annotated and truncated start codons associated with a rare disease (Supplemental Table S5).

**Figure 6.**
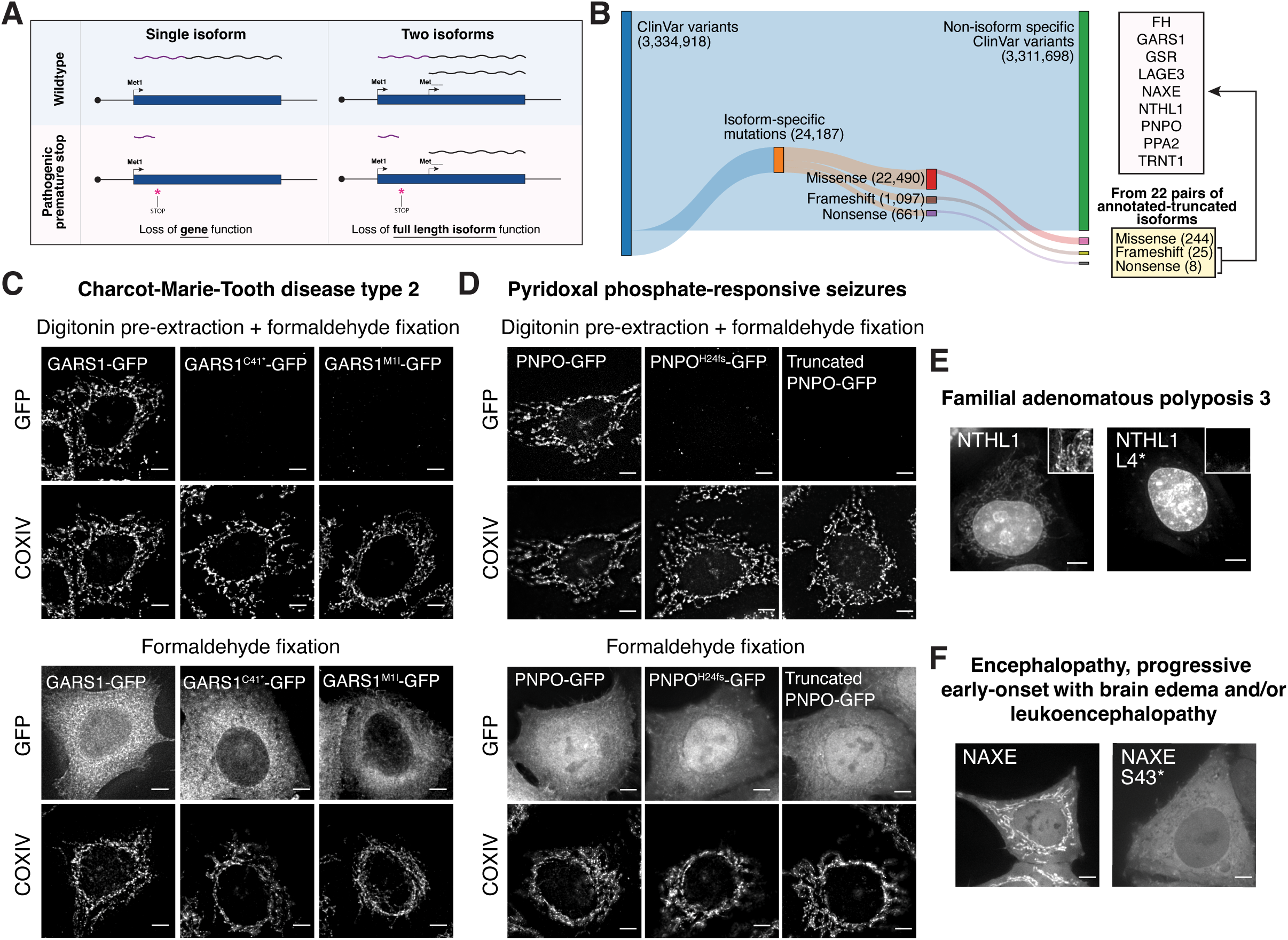
Rare disease mutations target specific alternative translational isoforms. (**A**) Schematic model showing how a pathogenic premature stop codon mutation would affect a gene with one (left) or multiple (right) translation initiation sites. With multiple alternative start codons, the premature stop codon mutation might not eliminate all protein variants from the gene. Similarly, frameshift or missense mutations may selectively impact one or multiple alternative translational isoforms. (**B**) Sankey plot summary of isoform specific mutations from the ClinVar database. From the 22 pairs of differentially localized annotated-truncated isoforms, 9 of these pairs had isoform specific mutations that would eliminate the protein product from the first start codon. (**C**) Immunofluorescence of GARS1 5′ UTR – CDS – GFP with different ClinVar mutations, co-stained with COXIV (mitochondrial marker) in cytosolic pre-extracted (top) or formaldehyde fixed cells (bottom). Wild type GARS1 localizes to the cytosol, nucleus, and mitochondria, whereas the C41* and M1I pathogenic variants eliminate the usage of the first start codon of GARS1 leading to only cytosolic/nuclear localization. (**D**) Similar as 6C except with PNPO instead of GARS1. (**E**) Live cell imaging of wild type of pathogenic L4* NTHL1 5′ 5′ UTR – CDS – GFP. Inset region zooming in on mitochondrial localization is indicated by white box. Inset images are scaled to be brighter than full size image. (**F**) Live cell imaging of wild type of pathogenic S43* NAXE 5′ 5′ UTR – CDS – GFP. Scale bar, 5 µm.

To test the impact of these mutations, we evaluated specific genes. We identified mutations in NTHL1 (c.11T>A (p.Leu4Ter) in Familial adenomatous polyposis 3, GARS1 (c.123C>A (p.Cys41Ter) in Charcot-Marie-Tooth disease type 2, and PNPO (c.69dup (p.His24fs) in patients with pyridoxal phosphate-responsive seizures, amongst others (Fig. 6B; Fig. S6B-E, Supplemental Table S5). These premature stop codons and frameshift mutations are typically interpreted as completely eliminating the production of a protein (Fig. 6A). However, the presence of a downstream alternative translation initiation sites for these genes instead predicts that these mutations would only eliminate the upstream protein isoform, leaving the alternative N-terminal protein isoform intact (Fig. 6A). Indeed, we found that these disease alleles selectively abolished the expression of mitochondrial isoforms initiated from the first AUG while preserving the cytosolic/nuclear-localized downstream isoform (Fig. 6C-E). Similarly, a previous study ^54^ identified patients with a c.128C>A (p.Ser43Ter) premature stop codon variant in NAD(P)H-hydrate epimerase (NAXE). These patients exhibit early-onset progressive encephalopathy characterized by brain edema and/or leukoencephalopathy. One such patient (LOVD ID: 00334932) was homozygous for the NAXE p.Ser43Ter variant, with the disease inherited in an autosomal recessive manner, suggesting that this is a loss-of-function allele. In contrast to wild type NAXE, which produces dual mitochondrial and cytosolic isoforms, we found that the NAXE Ser43Ter variant only produces a cytosolic/nuclear isoform of NAXE, suggesting that this disease is caused by the specific loss of the mitochondrial NAXE isoform (Fig. 6F).

These results highlight the significance of alternative start codon usage and N-terminal isoforms in the context of rare diseases, emphasizing the need to consider isoform-specific impacts of mutations in genetic and clinical studies. To facilitate the identification of isoform-specific mutations for newly generated patient genetic sequencing data, we developed SwissIsoform—a platform designed to determine whether a mutation selectively affects one or more protein isoforms for genes with a translational truncation (Fig. S6F).

### TRNT1 isoform-selective mutations in atypical SIFD patients

We next sought to leverage the understanding of dual-localized isoforms to evaluate whether isoform-specific mutations can improve our understanding of rare disease pathologies. In a cohort of patients with sideroblastic anemia, B-cell immunodeficiency, periodic fevers, and developmental delays (SIFD) at Boston Children’s Hospital, we identified several isoform-specific mutations in TRNT1 (Fig. 7A). In patients with canonical SIFD, mutations downstream of methionine 30 (Met30)—including premature stop codons, frameshifts, and catalytic region mutations—are predicted to disrupt both the mitochondrial and nuclear TRNT1 isoforms ^55^. Interestingly, amongst the cohort of Boston Children’s Hospital SIFD patients, we identified two patients with unique SIFD phenotypes that carry mutations predicted to selectively impact specific TRNT1 isoforms (Fig. 7B).

**Figure 7.**
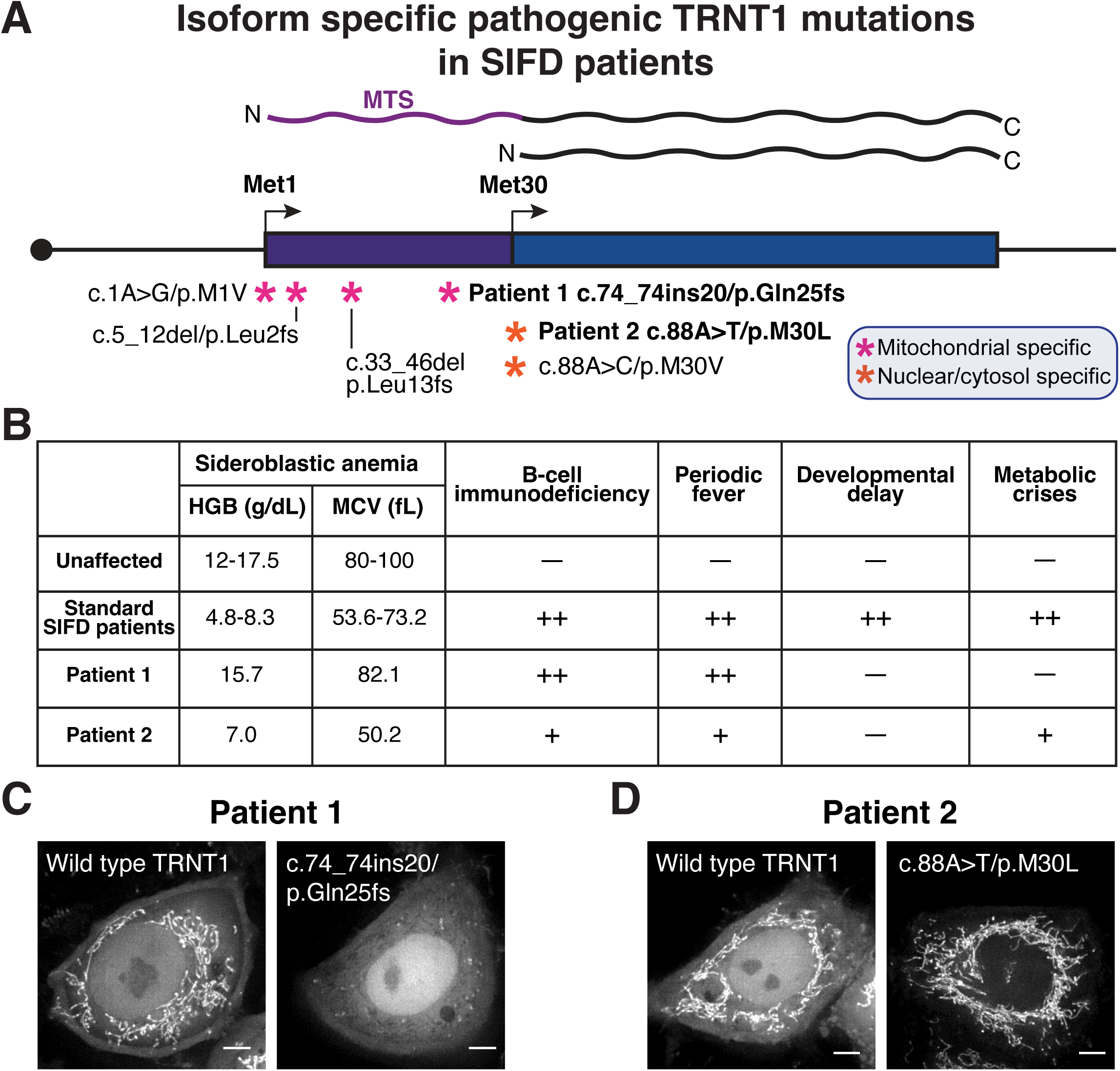
TRNT1 isoform-selective mutations in atypical SIFD patients. (**A**) Schematic representation of isoform specific TRNT1 SIFD mutations from ClinVar and patients from Boston Children’s Hospital. (**B**) Table showing severity of clinical symptoms from SIFD patients. HGB and MCV measurements for standard SIFD patients from ^56^ and unaffected individuals from ^61^. HGB = hemoglobin levels, MCV = mean corpus volume. Metabolic crises include nausea, vomiting, diarrhea, and abdominal pain. (**C**) Live cell imaging of wild type and patient 1 TRNT1 isoform specific mutation. (**D**) Live cell imaging of wild type and patient 2 TRNT1 isoform specific mutation. Scale bar, 5 µm.

Patient 1 is a developmentally normal 58-year-old male with a lifelong history of periodic fevers treated since childhood with colchicine and/or TNFαinhibitors for a presumptive diagnosis of Familial Mediterranean Fever (FMF). This patient has received 5 transfusions during his lifetime in the context of recurrent pulmonary and gastrointestinal hemorrhage. However, in contrast to classic SIFD patients, he is not chronically anemic. The patient was diagnosed with chronic myeloid leukemia at age 42 and treated into molecular remission with dasatinib, and subsequently developed total color and night blindness. In recent years, he was noted to be hypogammaglobulinemic, and the fevers have accelerated in frequency and associated with multiple episodes of culture-negative bronchitis and a pustular rash. Molecular analysis for periodic fevers at age 50 revealed TRNT1 mutations (NM_182916.3, c.74_74ins20/c.1056+11G; p.Gln25His*6/p.?) confirmed to be biallelic by segregation analysis in his parents and a brother (Fig. S7A). Patient 1 is not anemic (HGB 15.7 g/dL) but is slightly microcytic (MCV 82.1 fL), and has not had a bone marrow examination (Fig. 7B). These clinical features are in contrast to the classic SIFD patient with mutations targeting both isoforms of TRNT1, who are severely anemic ^56^. Although canonical gene models predict that the p.Gln25His*6 mutation would be a null allele, we found that expression of a TRNT1-GFP construct containing the patient frameshift mutation selectively eliminated the mitochondrial isoform but retained nuclear localization (Fig. 7C), as would be predicted based on our leaky scanning model for TRNT1 decoding. We also identified another patient from ClinVar (SCV005069659.1) with reported retinal dystrophy without anemia similar to patient 1 and a mitochondrial TRNT1 isoform-specific mutation (c.33_46del; p.Leu13fs) (Fig. S7B). These findings suggest that the unique SIFD clinical phenotypes in patient 1 may arise from the selective disruption of mitochondrial TRNT1, rather than disrupting all TRNT1 isoforms.

We also identified a second patient with isoform-specific TRNT1 mutations and distinct SIFD symptoms. Patient 2 is a 10-year-old developmentally normal Haitian/African American child who presented at age 16 months with microcytic anemia (HGB 7.0 g/dL, MCV 50.2 fL) and oral candidiasis, mild B cell and CD8+ T cell lymphopenia without hypogammaglobulinemia (Fig. 7B). Bone marrow examination showed numerous ringed sideroblasts and genetic testing revealed biallelic TRNT1 mutations (c.88A>T/c.461C>T; p.Met30Leu/p.Thr154Ile) (Fig. S7C). Ophthalmologic examination showed mild retinal dystrophy. His subsequent clinical course has been characterized by a stable anemia and 2-3 febrile episodes associated with abdominal pain, nausea, and vomiting per year that typically respond to hydration alone. Of the compound heterozygous mutations identified in Patient 2, Thr154Ile occurs in the catalytic region and is predicted to alter the function of both nuclear and mitochondrial TRNT1 isoforms (AlphaMissense score = 0.989 ^57^). In contrast, the M30L mutation impacts the alternative downstream start codon mutation in TRNT1 and is predicted to disrupt translation initiation at the alternative TRNT1 start codon, reducing the nuclear isoform. Consistent with this, C-terminal GFP fusion assays showed a strong depletion of nuclear TRNT1, while the mitochondrial isoform remained largely unaffected (Fig. 7D). In addition to patient 2, a previously reported Chinese patient carrying a c.88A>G/ c.363G > T allele (p.Met30Val/p.Glu121Asp) also had a similarly mild phenotype characterized by normal neurocognitive development and recurrent pulmonary infections and autoimmune musculoskeletal complications ^58^. Modeling this TRNT1 M30V allele also disrupted the nuclear truncated TRNT1 isoform, leaving the mitochondrial TRNT1 isoform unaffected (Fig. S7D-E). These results suggest the unique SIFD phenotype in patient 2 arises from the loss of nuclear TRNT1 activity, with mitochondrial function preserved.

These findings highlight the importance of considering alternative translation initiation in interpreting patient mutations. Without accounting for TRNT1 alternative initiation, mutations like Gln25fs would be misclassified as complete loss-of-function mutations. Similarly, the Met30Leu and Met30Val mutation is predicted to be benign using tools like AlphaMissense (score = 0.278/1.0 and 0.135/1.0 ^57^), but should be considered pathogenic due to its impact on alternative translation initiation. Finally, our findings suggest that the diverse and multifaceted phenotypes observed in SIFD are likely driven by the dual roles of TRNT1 in both mitochondrial and nuclear functions.

## Discussion

Mitochondrial endosymbiosis required the ability to perform the same core biological activities both within the mitochondria and within the host cell ^6^. One key strategy for adapting to mitochondrial acquisition involves paralog duplication/gene transfer, followed by the evolution of a mitochondrial targeting sequence in one paralog to enable dual functionality in both compartments. Here, we demonstrate that alternative start codon selection provides an additional mechanism for generating dual-localized mitochondrial and nuclear/cytosolic protein isoforms, ensuring the maintenance of core cellular processes across these distinct locations (Fig. 2). Furthermore, we provide evidence that alternative start codon usage has early eukaryotic origins, potentially arising from a bacterial/archaebacterial ancestor soon after the emergence of mitochondria (Fig. 3). Beyond dual-localized mitochondrial/cytosolic proteins, we also identify alternative protein variants that localize to other organelles, suggesting a broader role for alternative translation initiation in organelle-specific proteome diversification.

To achieve the dual production of protein isoforms, alternative start codon selection is influenced by multiple cis-elements, including leaky ribosome scanning through weak start codons, RNA secondary structure, uORF reinitiation, alternative promoter usage, and alternative splicing. Our work also revealed a critical role of 5′ UTR length in modulating start codon selection (Fig. 4), with implications for how alternative promoter usage and 5′ UTR disease variants impact N-terminal isoform production. For instance, upstream transcription initiation through the usage of an alternative promoter or a disease-associated 5′ UTR insertion in a gene with a normally short 5′ UTR could extend its length, favoring the first AUG start codon over downstream sites. Our findings also underscore key considerations for studying translation in mRNAs with short 5′ UTRs, including the choice of transgenic promoters, which can extend the 5′ UTR and disrupt isoform production (see Fig. 4A and S4A).

The use of alternative start codons within a single mRNA also has important implications for interpreting disease mutations (Fig. 6 and 7). Rather than assuming a premature stop codon or frameshift mutation eliminates all protein function, the presence of alternative start codons may instead disrupt the production of specific isoforms. We show that nonsense or frameshift mutations between two translation initiation sites selectively eliminate the longer protein isoform while having limited effects on downstream initiation. Notably, we highlight isoform-specific TRNT1 mutations as key factors in the heterogeneous clinical phenotypes of SIFD (Fig. 7). Beyond premature stop and frameshift mutations, missense mutations that affect specific isoforms may also contribute to disease. Mutations that alter the Kozak context of an upstream start codon, annotated start site, or downstream truncation could also lead to the altered expression of alternate isoforms. Furthermore, mutations affecting alternative promoters or RNA secondary structure may similarly be pathogenic by altering protein isoform ratios. Given the plethora of rare disease with unexplored molecular mechanisms and variants of unknown significance ^59^, our findings underscore the need to systematically identify and analyze rare disease mutations that impact specific protein isoforms.

## Acknowledgments

This work was supported by grants from the NIH (R35GM126930 to I.M.C.; R01DK087992 and R24DK094746 to M.F.; and AI144369 and AI158501 to S.L), and the Chan Zuckerberg Initiative Rare as One Project grant to I.M.C. J.L. and Y.T. are supported in part by the Natural Sciences and Engineering Research Council of Canada. M.D. is supported in part by a NSF GRFP fellowship. We thank the Whitehead Genome Technology Core for sequencing; the Whitehead Quantitative Proteomics Core for mass spectrometry; the Whitehead Flow Cytometry Core for cell sorting; Brittania Moodie for technical assistance, Glenn Li (Bartel lab) for providing *S. cerevisiae* RNA; Alexandra Wilson, David Bartel, Heather Keys, and all members of the Cheeseman and Bartel labs for helpful discussions; Olivia Rissland for constructive comments on the manuscript; the Bartel lab for use of equipment and reagents; and Lyra Cheeseman for conducting initial predictions of protein localization.

## Author contributions

J.L. and I.M.C. conceived and designed experiments; All experiments and analysis were performed by J.L., except for 5′ RACE and sequencing by Y.T., ClinVar analysis by M.D, assisted with reporter assays E.K., *T. gondii* TRNT1 imaging by C.J.G., and M.D.F. for clinical observations; J.L. and I.M.C. wrote the manuscript with input from all authors; I.M.C, S.L., M.D.F., J.L., Y.T., M.D. acquired funding; I.M.C. and S.L. supervised the project.

## Data and materials availability

The mass spectrometry proteomics data have been deposited to the ProteomeXchange Consortium via the PRIDE ^60^ partner repository with the dataset identifier PXD062112 and 10.6019/PXD062112. Code is available at Github (https://github.com/cheeseman-lab/swissisoform).

---

**Supplementary figure 1.**
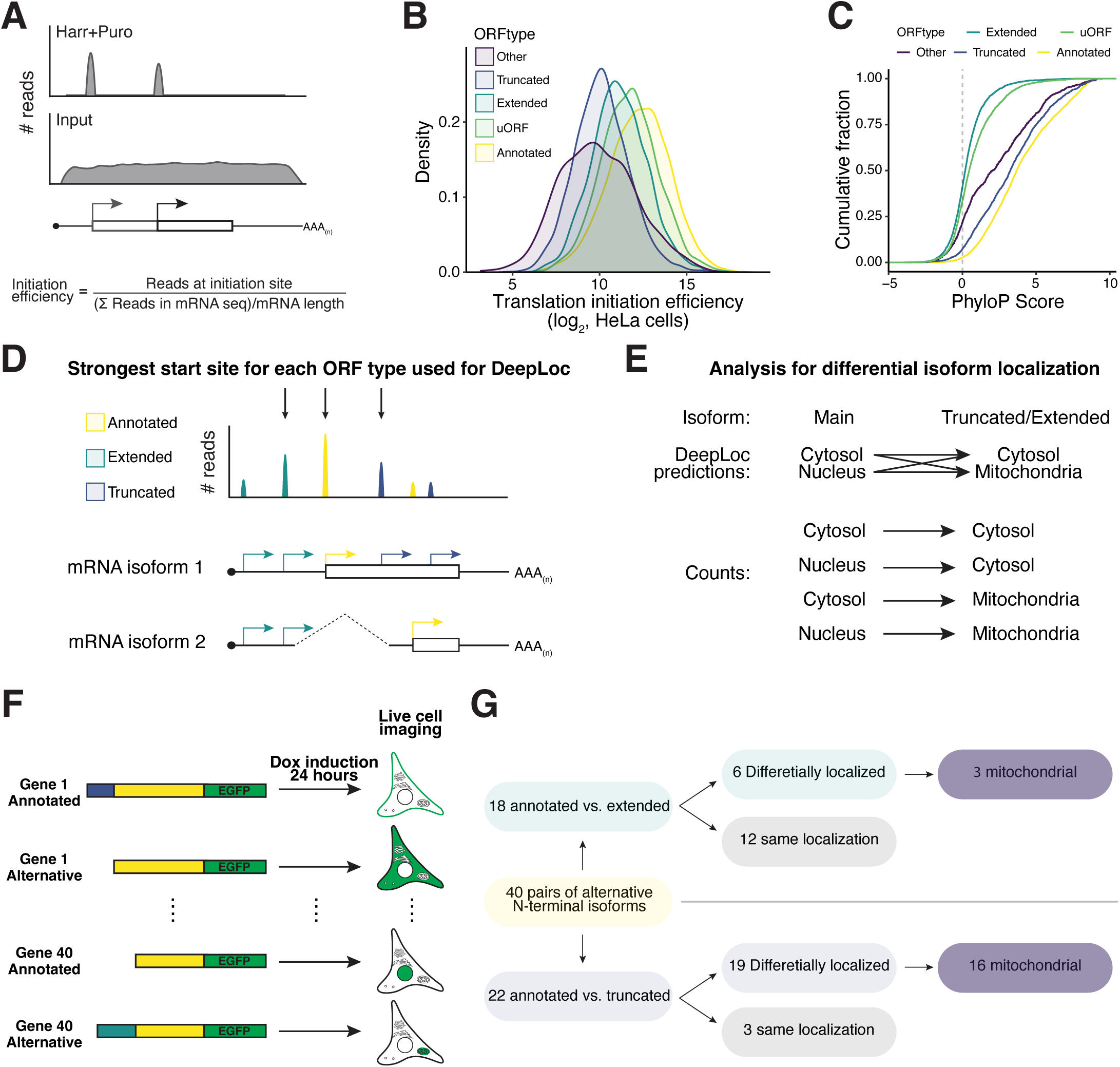
Alternative translation initiation drives distinct subcellular localizations. (**A**) Schematic representation of translation initiation site efficiency measurement. (**B**) Histogram showing relative translation initiation efficiency between ORF types from asynchronous HeLa cells ^33^. (**C**) CDF plot showing the average PhyloP score of start codons for each ORF types. (**D**) Schematic showing the selection of N-terminal isoform for each ORF type if a gene has multiple translation initiation sites. The start site with the strongest translation initiation efficiency is selected and used for DeepLoc analysis in Figure 1. (**E**) Schematic showing the analysis of DeepLoc output. If the annotated isoform is predicted mitochondrial and the truncated isoform is predicted cytosolic and nuclear, the predicted change for the associated gene is mitochondria to cytosol and mitochondria to nucleus. Additional details described in the Methods. (**F**) Schematic representation of localization screen (40 alternative N-terminal isoforms) to assess the localization of selected alternative N-terminal isoforms. (**G**) Summary of the results from localization screen.

**Supplementary figure 2.**
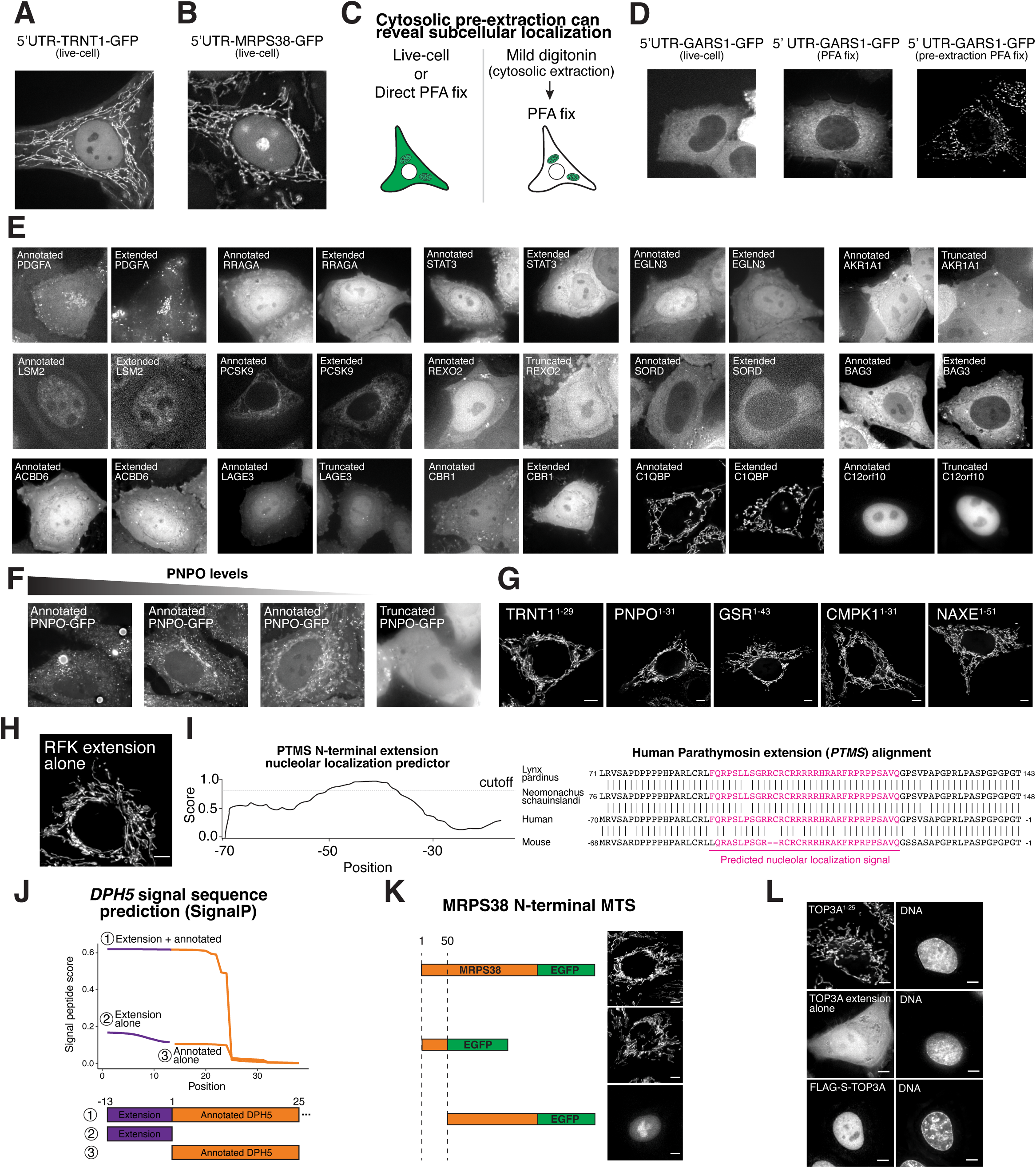
Mechanisms underlying differential localization for alternative N-terminal isoforms. (**A**) Live cell imaging of TRNT1 5′ UTR – CDS – GFP shows dual localization to nucleus/cytosol and mitochondria. (**B**) Live cell imaging of MRPS38/AURKAIP1 5′ UTR – CDS – GFP shows dual localization to nucleolus and mitochondria. (**C**) Schematic showing the possible differences between live-cell or direct formaldehyde fixed and cytosolic pre-extracted imaging. Pre-extraction of the cytosolic signal may uncover subcellular localization that is being obscured by the cytosolic signal. (**D**) Localization of GARS1 5′ UTR – CDS – GFP by live-imaging, direct formaldehyde fix, and pre-extraction followed by formaldehyde fixed immunofluorescence. Since the cytosolic isoform of GARS1 is more highly expressed than the mitochondrial isoform, the mitochondrial localization of GARS1 can by clearly observed by pre-extracting cytosolic signal. (**E**) Live cell imaging of selected alternative translational isoforms where the same localization was observed. (**F**) An example where the localization of an isoform may be altered based on expression level. Cells with high expression of the PNPO Met1 isoform exhibit protein aggregation into foci at high concentrations, whereas low concentrations lead to mitochondrial localization. (**G**) Live cell imaging showing the localization of the N-terminal region tagged with C-terminal GFP of selected genes before the truncation start codon. (**H**) Live cell imaging of the RFK extension alone (lacks the annotated protein sequence) tagged with C-terminal GFP. (**I**) Left, nucleolar localization sequence detector ^62^ output for Human PTMS N-terminal extension. Right, sequence alignment of annotated PTMS from lynx, seal and the N-terminal extension of PTMS from human and mouse. Highlighted in pink is the predicted nucleolar localization signal based on Nucleolar localization sequence detector. (**J**) SignalP prediction ^63^ for signal sequences for the indicated DPH5 sequences. Combining the N-terminal extension of DPH5 with the N-terminus of DPH5 creates a signal sequence as shown in Fig. 2D. (**K**) Live cell imaging of the indicated MRPS38/AURKAIP1 construct. (**L**) Live cell imaging of cells expressing the indicated TOP3A constructs. Scale bar, 5 µm.

**Supplementary figure 3.**
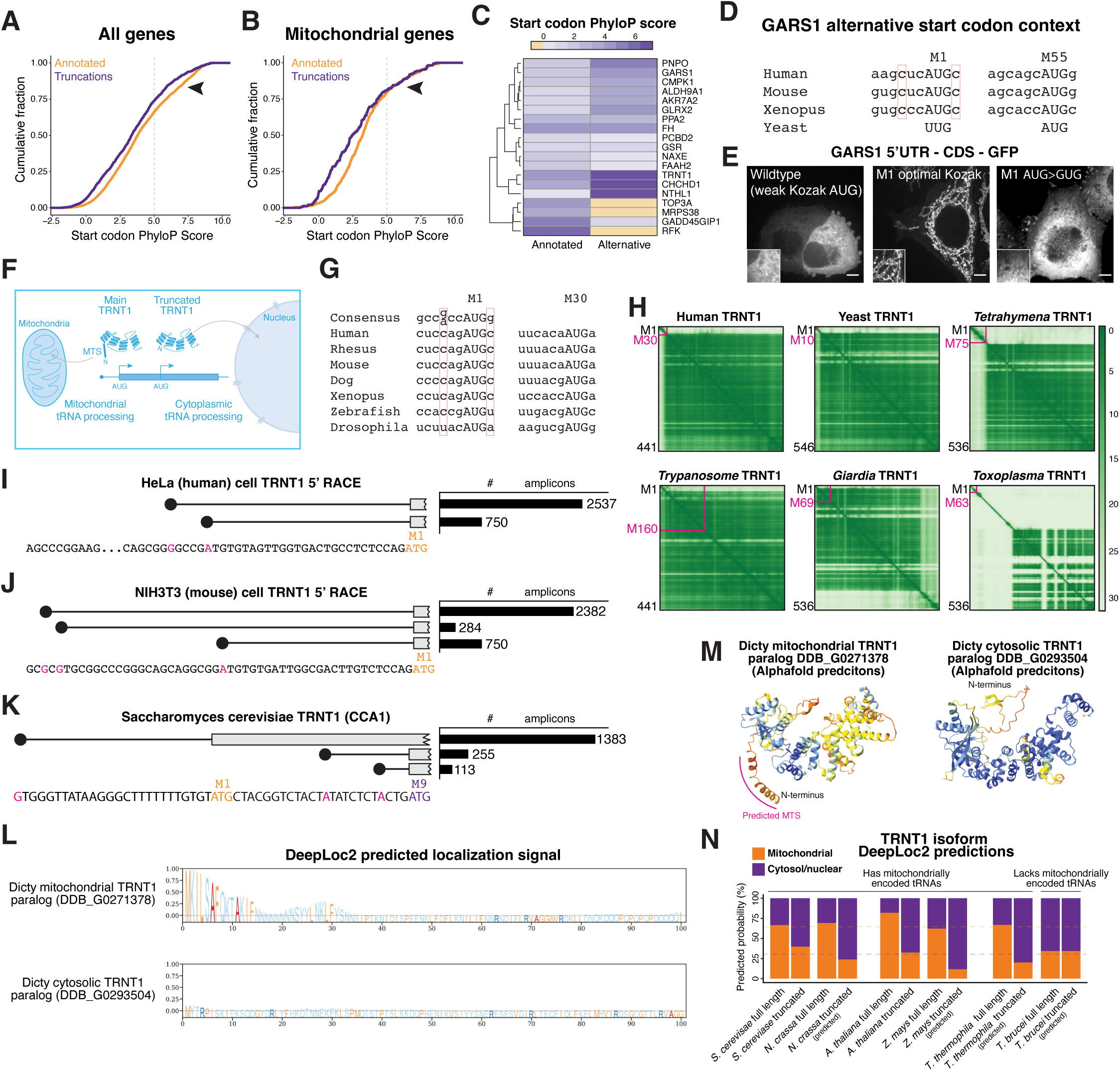
Evolutionary analysis of alternative N-terminal isoforms. (**A**) Cumulative distribution function plot showing the average PhyloP score (vertebrate 100 way) for all annotated and truncated ORFs. (**B**) Same as (A) but only for mitochondrial genes using annotations from MitoCarta ^64^. (**C**) Heatmap showing the start codon PhyloP score for annotated and alternative ORFs that displayed differential mitochondrial localization. (**D**) Sequence alignment from listed organisms around the GARS1 start codons. The weak Kozak context ^65^ of GARS1 at the first AUG to allow for leaky ribosome scanning is highly conserved with budding yeast having a non-AUG start codon instead of a weak AUG. (**E**) Live cell imaging showing the localization of GARS1 5′ UTR – CDS – GFP with the indicated mutations to the first start codon. Scale bar, 5 µm. (**F**) Schematic model for the role of TRNT1 alternative start codons in producing differentially localized protein isoforms to satisfy tRNA biogenesis for nuclear and mitochondrially-encoded tRNAs. (**G**) Sequence alignment around the start codons of TRNT1. Consensus sequence is for Vertebrate organisms ^65^. The red box is highlighting the −3 and +4 position–particularly important positions for the Kozak context. (**H**) Alphafold predicted aligned error plot for TRNT1 in the indicated organisms. The magenta line indicates the position of the second in-frame AUG start codon coding for the N-terminal nuclear TRNT1 isoform. For Humans, Yeast, and Tetrahymena, the N-terminal truncation removes an unstructured region. For *T. brucei* and *G. lambia*, the predicted N-terminal truncation would remove a large part of the protein’s catalytic domain. *T. gondii* TRNT1 N-terminal region is very long and unstructured without and mitochondrial prediction signals. Scale bar indicates expected position error (Ångstrohm). (**I**) 5′ RACE and sequencing from TRNT1 in HeLa (human). The TRNT1 5′ UTR for ensembl ENST00000251607 transcript is shown. Not shown are small number of reads where the 5′ end maps after Met30, which likely represents degradation product or translates to a protein that removes most of the catalytic domain. The magenta nucleotide represents the empirically determined 5′ end in HeLa cells. (**J**) Same as (I) except in mouse NIH3T3 cells. The TRNT1 5′ UTR for ensembl ENSMUST00000113248 transcript is shown. (**K**) 5′ RACE and sequencing from TRNT1 in *S. cerevisiae*. (**L**) DeepLoc2.1 prediction of the two *D. discoideum* TRNT1 paralogs. Note that the next in-frame AUG for the mitochondrial TRNT1 paralog (DDB_G0271378) is Met268, suggesting that encoding two isoforms in this gene is unlikely. Hypothetical usage of this downstream AUG would remove most of the TRNT1 catalytic domain, rendering the protein inactive. (**M**) Alphafold structure predictions for the two *D. discoideum* TRNT1 paralogs, colored by model confidence. (**N**) DeepLoc2.1 predictions for TRNT1 isoforms from the indicated organism. For organisms where there is no experimental evidence for truncated TRNT1 isoform, the second in-frame methionine was used. Dashed line indicates average predicted mitochondrial localization probability for *Saccharomyces cerevisae*, *Neurospora crassa*, *Arabidopsis thaliana*, *Zea mays*, and *Tetrahymena thermophila* annotated TRNT1 (orange) or truncated TRNT1 (purple). Similar to *T. gondii* (Fig. 2E and 2J), *T. Brucei* lack mitochondrial tRNAs ^39^ and DeepLoc predicts only nuclear/cytosolic TRNT1 localization.

**Supplementary figure 4.**
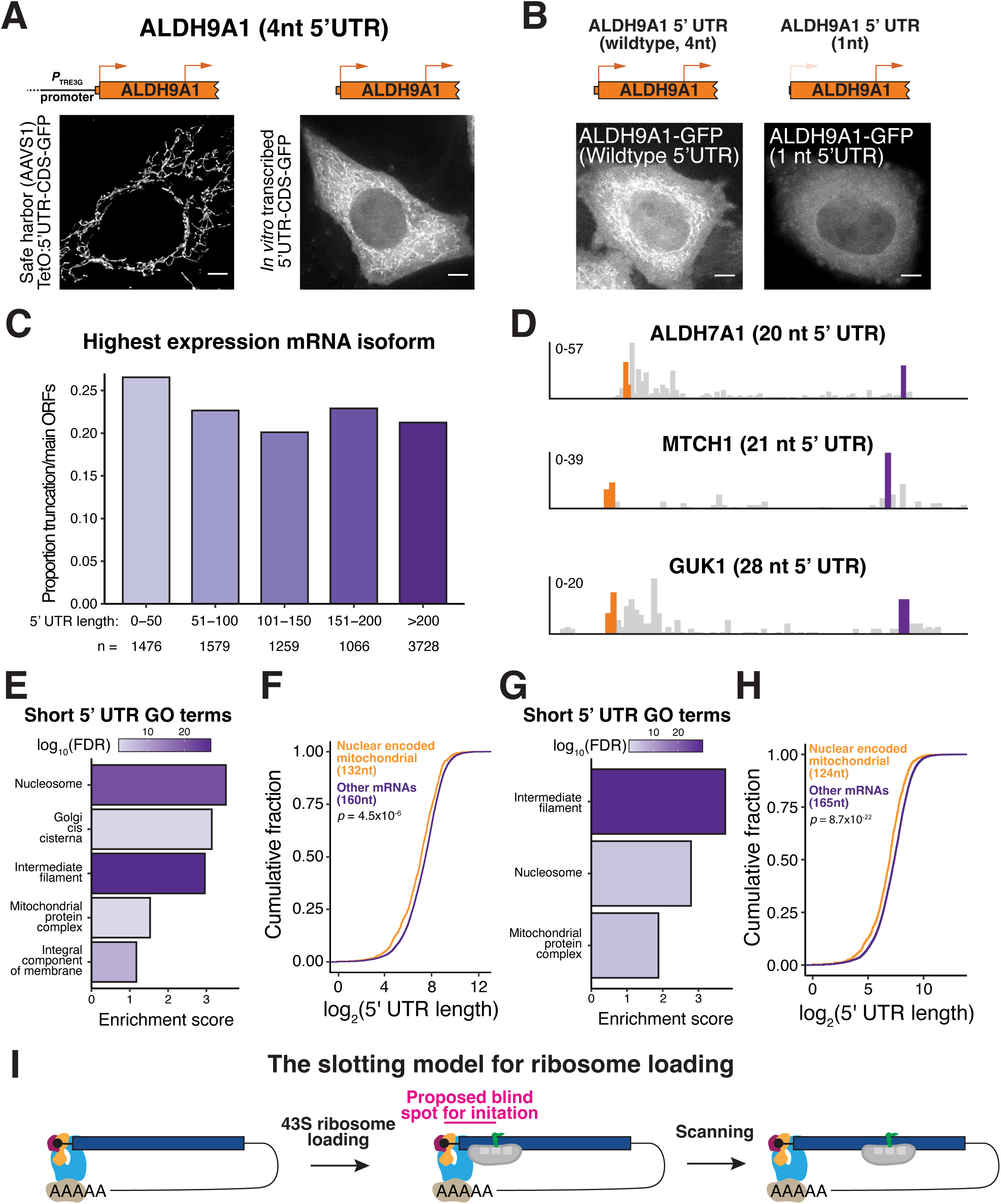
Additional analysis of 5′ UTR lengths and alternative N-terminal isoform expression. (**A**) Live cell imaging of ALDH9A1 5′ UTR – CDS – GFP expressed under a dox-inducible promoter (left) or through transfection of an *in vitro* transcribed mRNA. (**B**) Live cell imaging of wild type (left, 4 nt) and short (right, 1 nt) ALDH9A1 5′ UTR – CDS – GFP. (**C**) Same as Fig. 4I except using the most highly expressed transcript from HeLa cells for each gene rather than the longest transcript isoform. (**D**) Ribosome profiling trace of selected genes that possibly undergo alternative start codon selection through their short 5′ UTR length. Orange bars represent annotated start codon reads whereas the purple bar indicates alternative start site reads. ALDH7A1, MTCH1, and GUK1 have 20, 21, and 28nt 5′ UTR, respectively. (**E**) GO-term enrichment analysis of short 5′ UTRs (≤50 nt compared to >50 nt 5′ UTRs) using the transcript isoform with the longest coding sequence length for each gene. (**F**) CDF plot of 5′ UTR lengths of nuclear-encoded mitochondrial genes and other nuclear-encoded genes using the transcript isoform with the longest coding sequence length for each gene. (**G**) Same as (E) except using the most highly expressed transcript from HeLa cells for each gene rather than the longest transcript isoform. (**H**) Same as (F) except using the most highly expressed transcript from HeLa cells for each gene rather than the longest transcript isoform. (**I**) Schematic model highlighting the proposed blind spot for start codon selection on mRNAs with short 5′ UTRs ^45^.

**Supplementary figure 5.**
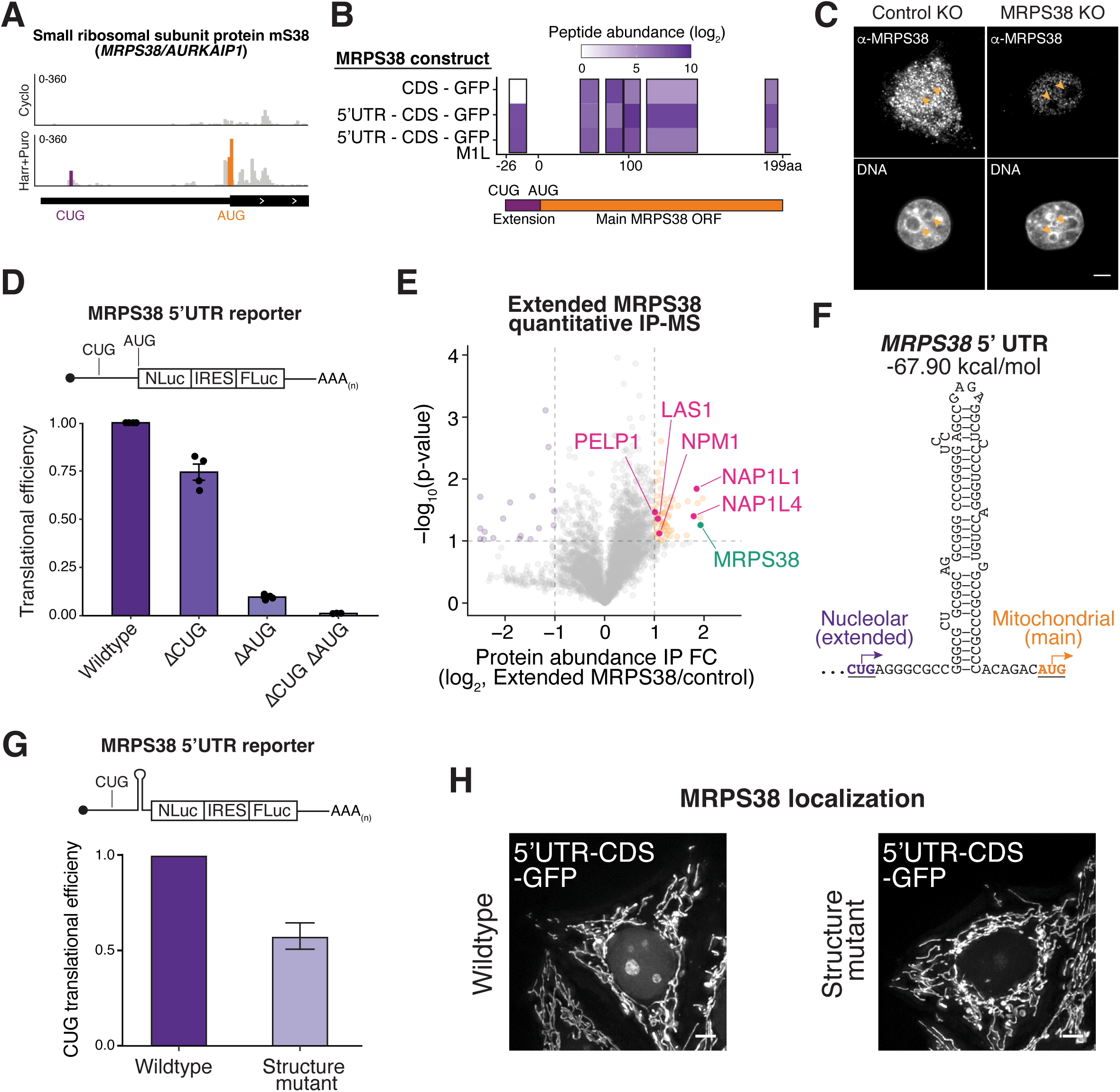
Validation, function, and mechanisms of MRPS38/AURKAIP1 N-terminal extension. (**A**) Ribosome profiling reads around the MRPS38/AURKAIP1 alternative start codons. (**B**) Mass spec peptide coverage across the MRPS38/AURKAIP1 N-terminal extension for the indicated MRPS38 immunoprecipitation. Constructs with the MRPS38 5′ UTR results in the detection of a peptide mapping to MRPS38 extension. (**C**) Immunofluorescence images of control and MRPS38 knockout cells (both isoform) stained with anti-MRPS38. Orange arrowhead indicates nucleolus. Scale bar, 5 µm. (**D**) Top, schematic of luciferase reporter used to assess MRPS38 start codon activities. Bar plot showing luciferase reporter assays with the MRPS38/AURKAIP1 5′ UTR. The indicated start codons were mutated. Mutation of the extended MRPS38 extension CUG start site reduces nanoLuc activity. IRES-initiated FLuc activity accounts for changes in mRNA abundance and transfection efficiency ^33^. (**E**) Volcano plot showing quantitative IP-MS analysis of extended MRPS38. Orange points represent proteins that are enriched in the extended MRPS38 IP. Pink points are selected nuclear/nucleolar proteins. N = 2 biological replicates. (**F**) RNA secondary structure downstream of the CUG MRPS38 start codon. (**G**) Top, luciferase reporter used to assess the efficiency of the CUG initiated MRPS38 extension start site. Bar plot showing the effect of mutating the RNA secondary structure on MRPS38 CUG translation. (**H**) Left, live cell imaging of wild type MRPS38/AURKAIP1 5′ UTR – CDS – GFP shows dual localization. Right, localization of MRPS38/AURKAIP1 5′ UTR – CDS – GFP with silent mutations that disrupt the RNA secondary structure reduces nucleolar localization (CUG initiated isoform). Scale bar, 5 µm.

**Supplementary figure 6.**
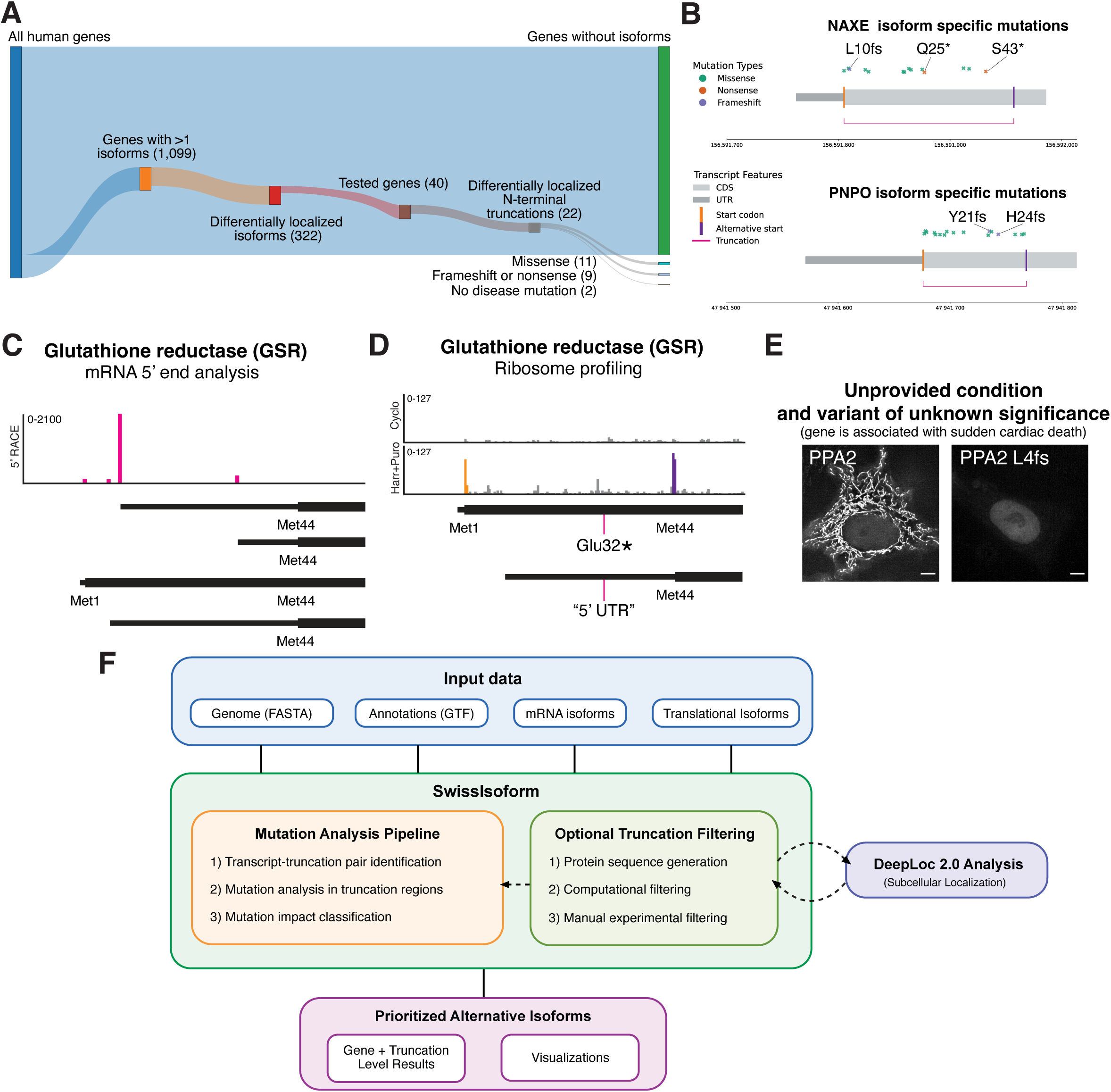
Pathogenic mutations target specific alternative N-terminal isoforms. (**A**) Sankey plot summarizing the number of genes with isoform specific ClinVar mutations. Of the 22 genes that produce differentially localized N-terminal truncations, 9 of them have frameshift or nonsense mutations that eliminate the annotated but keep the truncated isoform intact. (**B**) Examples of genes with isoform specific ClinVar mutations. (**C**) Bar graph showing the number of 5′ end reads from GSR 5′ RACE sequencing showing, suggesting that the alternative GSR isoforms are regulated transcriptionally. (**D**) Ribosome profiling trace around GSR start codons, indicating a pathogenic mutation that is predicted to eliminate full length but not truncated GSR protein isoform. (**E**) Live cell imaging of wild type of variant of unknown significance L4fs PPA2 5′ 5′ UTR – CDS – GFP. Transfected *in vitro* transcribed reporters were used for this analysis given the short 5′ UTR (12 nt) of PPA2. Scale bar, 5 µm. (**F**) Schematic outline of Swissisoform, a platform to identify patient mutations that specifically target an alternative translational isoform.

**Supplementary figure 7.**
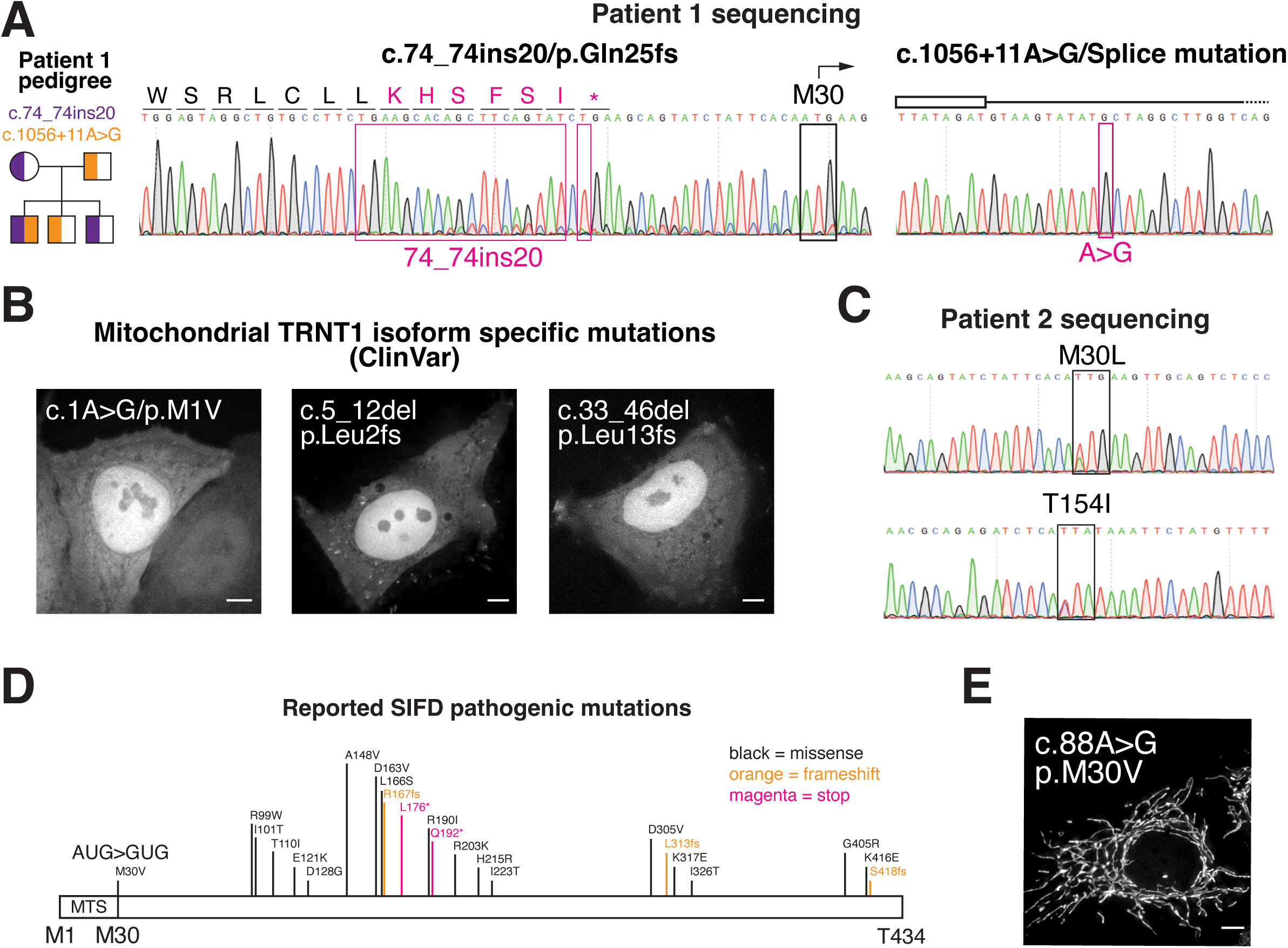
TRNT1 mutations in SIFD patients. (**A**) Pedigree and TRNT1 sequencing data from patient 1. TRNT1 from patient 1 blood samples were subcloned and the Sanger sequencing trace for resulting clones are displayed. (**B**) Live-cell imaging of TRNT1 5′ UTR – CDS – GFP with the indicated patient mutation. (**C**) Sanger sequencing trace for TRNT1 from patient 2 blood samples. (**D**) Reported pathogenic SIFD mutations in TRNT1 from ^66^, revealing another truncated TRNT1 specific mutation. (**E**) Live cell imaging of TRNT1 5′ UTR – CDS – GFP with the indicated patient mutation. Scale bar, 5 µm.

**Supplemental table 1. DeepLoc predictions for alternative N-terminal isoforms.**

**Supplemental table 2. Experimentally determined subcellular localization for 40 alternative N-terminal isoforms.**

**Supplemental table 3. Genes with short 5′ UTRs and multiple start codons**

**Supplemental table 4. Quantitative extended MRPS38/AURKAIP1 IP-MS analysis.**

**Supplemental table 5. Isoform specific ClinVar analysis**

## Experimental Procedures

### Ethics approval

Human samples were obtained with written informed consent under a human subject’s research protocol (06-12-0536) approved by the institutional review board at Boston Children’s Hospital.

### Tissue culture

HeLa and HEK293T cells were cultured in DMEM with 10% heat-inactivated fetal bovine serum, 2 mM L-glutamine and 100 U/mL penicillin-streptomycin at 37°C with 5% CO2.

### Molecular biology and cell line generation

All cDNA, except for non-human genes analyzed in this study, was amplified from HeLa cell cDNA. HeLa cell cDNA was generated via reverse transcription using the Maxima First Strand cDNA Synthesis Kit (K1641), following the manufacturer’s instructions. A summary of the protein sequences tested can be found in Supplemental table S2. Non-human genes were synthesized by Twist Bioscience.

To test whether the N-terminus contains sufficient signal sequences, we used the entire region upstream of the truncated start codon. For example, in the case of NAXE, where the N-terminal truncation occurs at Met52, we tested the region from Met1 to Val51 for a localization signal. For the analysis of N-terminal extensions, we included the entire extension, except in the case of DPH5. To determine whether extensions “mask” or “internalize” annotated N-terminal signal sequences, we appended a FLAG-S tag (MDYKDDDDKGKETAAAKFERQHMDSGGT) to the N-terminus of the annotated protein.

The amplified cDNA was inserted downstream of the TRE3G promoter and upstream of the MYC-TEV-EGFP tag using Gibson assembly. A SalI cut site was positioned between the TRE3G promoter and the cDNA construct. The plasmid also contained a puromycin resistance marker, a reverse tetracycline-controlled transactivator, and a C-terminally EGFP-tagged transgene, flanked by safe harbor AAVS donor homology arms. All transgenes were expressed from the safe harbor AAVS1 locus ^43^. Donor plasmid also contained the puromycin resistance marker, reverse tetracycline-controlled transactivator, the transgene, flanked by safe harbor AAVS donor homology arms. To generate dox-inducible GFP cell lines, 500 ng donor plasmid and 500 ng pX330-sgRNA AAVS1 were co-transfected using Lipofectamine 2000. 2 days post transfection, cells were selected with 0.4 µg/mL puromycin for at least 3 days.

When optimizing the Kozak sequence, the 6 nucleotides upstream of the start codon was mutated to GCCACC. The +4 position was kept as the endogenous sequence to not change the protein coding sequence. ^65^

To change the 5′ UTR length, parts of the *Xenopus* Beta Globin 5′ UTR (5′ – CTTGTTCTTTTTGCAGAAGCTCAGAATAAACGCTCAACTTTGG – 3’) were appended to the endogenous 5′ UTR sequence to achieve the intended length. When shortening the 5′ UTR length, the endogenous sequence is used, truncating the 5′ end to reach the intended length.

For MRPS38/AURKAIP1 the RNA secondary structure sequence was mutated from 5′-ttgagccgcgtcgaggtcgggcttgggaagggtcagcgggaggcCTGagGgcGccGggCgctgcGgcaggCggGccCggGgtCcaGccGagaggCtcAcctggGacCtgtggCcgCcgCccacaGacC-3’ to 5′-ttgagccgcgtcgaggtcgggcttgggaagggtcagcgggaggcCTGagAgcAccAggTgctgcAgcaggTggTccAggTgtTcaAccAagaggTtcAcctggTacTtgtggTcgTcgTccacaAacT-3’.

### Subcellular localization predictions

DeepLoc2.1 ^34^, PSORTII ^67^, SignalP ^63^, TargetP ^68^, and NoD ^62^ were used to predict subcellular protein localizations. For all predictions, we used the webserver and default settings to generate the predicted localizations.

We selected the strongest translation initiation site for each ORF type (extension, annotated, truncation) per gene prior to any localization predictions. To do this, we selected initiation sites with the highest translation initiation efficiency (ribosome footprint reads at the initiation site from harringtonine ribosome profiling samples normalized to mRNA abundance) from asynchronous HeLa cells ^33^.

To quantify localization changes and generate the Sankey plot, we compared the predicted localization of the annotated isoform with that of the alternative isoform. If an alternative isoform had multiple predicted localizations, each distinct transition was counted separately. For example, if the annotated isoform was predicted to localize to the mitochondria, whereas the alternative isoform localized to both the cytoplasm and nucleus, we recorded two transitions: (1) mitochondria to cytoplasm and (2) mitochondria to nucleus. Similarly, if the annotated isoform was cytosolic and nuclear, and the alternative isoform was mitochondrial, we classified this as (1) cytosol to mitochondria and (2) nucleus to mitochondria. When both isoforms had multiple localizations, all possible transitions were counted. For instance, if the annotated isoform was mitochondrial and nuclear, while the alternative isoform was cytosolic and nuclear, we recorded: (1) mitochondria to cytosol, (2) mitochondria to nucleus, (3) nucleus to cytosol, and (4) nucleus to nucleus.

For isoforms that shared the same localization we recorded no transitions across organelles. For example, if both isoforms localized cytoplasm and nucleus then we recorded 1) cytosol to cytosol and 2) nucleus to nucleus.

For the analysis of TRNT1, the sequences from Ensembl (human ENST00000251607; rhesus monkey ENSMFAT00000093651; mouse ENSMUST00000057578; frog ENSXETT00000066167; fish ENSDART00000113706; fly FBtr0301574) was used in the predictions. The current WBcel235 annotation has the “truncated” TRNT1 isoform as “annotated” due to a misannotation (hpo-31; F55B12.4.1). For our predictions, we identified the longest open reading frame and designated it as the “annotated” ORF.

When plotting predicted probability of localization using DeepLoc2.1 (Fig. 3J, Fig. 4F; Fig. S3N), we took the average predicted probability for nucleus and cytosol prior to plotting. This is because by default (for example GFP) most proteins are cytosolic/nuclear.

### Live cell imaging of dox-inducible GFP cell lines

The GFP transgene was induced by addition of 1 µg/mL dox for 24 hours prior to imaging. Cells on glass bottom plates were incubated in 0.1 μg/mL Hoescht for >30 minutes. Images were taken on the Deltavision Ultra (Cytiva) system using a 60x/1.42NA objective. 8 μm images were taken with z-sections of 0.2 μm. All images except for mitochondrial translation assays presented were deconvolved and max projected. When quantitatively comparing intensities between images, the images were scaled equivalently and raw images were used.

### Immunofluorescence

HeLa cells were seeded on poly-L-lysine coated coverslips and fixed in PBS + 4% formaldehyde at room temperature for 15 minutes. To pre-extract cytosolic signal the plate of cells were placed on ice and incubated with 10 mM HEPES, 10 mM NaCl, 5 mM KCl, 300 mM sucrose, and 0.015% digitonin for 2 minutes. Cells were then washed with 10 mM HEPES, 10 mM NaCl, 5 mM KCl, 300 mM sucrose then fixed with PBS + 4% formaldehyde at room temperature for 15 minutes. Following fixation, cells were washed 3 times with PBS + 0.1% Triton X-100 and blocked in AbDil for 30 minutes - 1 hour. Following blocking, cells were stained at room temperature for 1 hour with ChromoTek GFP-Booster Alexa Fluor 488 (1:1000, Proteintech, gb2AF488) and α-COXIV (1:1000, Cell Signaling, 119675S) diluted in AbDil. For MRPS38/AURKAIP1 immunofluorescence, cells were incubated with anti-AURKAIP1/MRPS38 (1:1000, Thermo, PA5-56869) overnight at 4°C. Cells were washed with PBS + 0.1% Triton X-100, 3 times then incubated in secondary antibody diluted in AbDil for 1 hour at room temperature. After secondary, cells were stained in 1 μg/mL Hoescht for 15 minutes at room temp, then washed 3 times with PBS + 0.1% Triton X-100 before mounting in PPDM (0.5% p-phenylenediamine and 20 mM Tris-Cl, pH 8.8, in 90% glycerol) and sealed with nail polish.

### Start codon conservation analysis

PhyloP scores from alignments of 99 vertebrates to human genome (100way) were used in analyses. For start codons, the average PhyloP score for the 3 nucleotides were used in analyses. MitoCarta3 ^64^ was used to define genes that are mitochondrial. We note that a change from a CUG to UUG or another non-AUG start codon between organisms may have a low PhyloP score but the usage of the codon may still be used across organisms.

### Endogenous Tagging of TGGT1_315810 (TRNT1) in *T. gondii*

Type I RH parasites were grown in human foreskin fibroblasts (HFFs, ATCC SCRC-1041) and maintained in DMEM (GIBCO) supplemented with 2 mM glutamine, 10% inactivated fetal bovine serum (IFS), and 10 µg/ml gentamicin. TGGT1_315810 was endogenously tagged in *T. gondii* using a previously described high-throughput (HiT) tagging system ^69^. Briefly, a cutting unit targeting the 3’ end of the coding sequence of TGGT1_315810 was cloned into a plasmid carrying a V5-TEV-mCherry-HA payload. Prior to transfection, 50 µg of this plasmid was linearized with BsaI-HF V2 (NEB) and dialyzed for 1 h. The linearized plasmid was co-transfected with a Cas9 expression vector into RHΔku80Δhxgprt parasites. During transfection, parasites were resuspended in transfection solution containing 0.15 mM CaCl2, 2 mM ATP, 5 mM glutathione, 15 - 60 µg of DNA, and cytomix (10 mM KPO4, 120 mM KCl, 5 mM MgCl2, 25 mM HEPES, 2 mM EDTA, 2 mM ATP, and 5 mM glutathione) up to 400 µl. Parasites were transfected by electroporation with two 150 µs pulses at 100 ms intervals at 1700 V. One day post transfection, parasites were selected with 25 µg/mL mycophenolic acid and 50 µg/mL xanthine pH-balanced with 0.3 N HCl. Correct integration of the construct was validated by PCR.

### Immunofluorescence of tagged TGGT1_315810

Parasites were infected onto a confluent monolayer of HFFs on coverslips. One day post infection, wells were washed with PBS and fixed with a 4% formaldehyde solution at room temperature for 20 min. Wells were washed 3x with PBS, permeabilized with 0.1% triton for 8 min, and washed again 3x with PBS. Wells were blocked for 10 min with 5% goat serum/5% fetal bovine serum in PBS. Wells were incubated with anti-HA (Biolegend #901501; 1:1000 dilution) and anti-HSP70 (a gift from Dominique Soldati; 1:1000 dilution) primary solution for 30 min at room temperature. Wells were then washed 3x with PBS, blocked, and stained with secondary solution (1:1000 anti-rabbit Alexa Fluor 594 ThermoFisher A11037, and 1:1000 anti-mouse Alexa Fluor 488 ThermoFisher A11029) for 30 min at room temperature. Wells were then washed 3x with PBS, dipped twice in dH2O, and mounted on glass slides with Prolong Diamond Antifade Mountant.

### Immunoprecipitations

Polyclonal GFP antibodies was coupled to Protein A beads as described in ^70^.

For MRPS38 GFP IP experiments to map N-terminal extension peptides, cells were washed in PBS and resuspended 1:1 in 50 mM HEPES pH7.4, 1 mM EGTA, 1 mM MgCl_2_, 100 mM KCl, 10% glycerol then flash frozen in liquid nitrogen. Cells were thawed after addition of an equal volume of 75 mM HEPES pH 7.4, 1.5 mM EGTA, 1.5 mM MgCl_2_, 150 mM KCl, 15% glycerol, 0.075% NP-40, 1X Complete EDTA-free protease inhibitor cocktail (Roche, 4693159001), and 1 mM PMSF. Cells were lysed by sonication and cleared by centrifugation at 21000x g for 30 minutes at 4°C. The supernatant was mixed with Protein A beads coupled to rabbit anti-GFP antibodies and rotated end-over-end at 4°C for 2 hours. Beads were rinsed twice in wash buffer (50 mM HEPES pH 7.4, 1 mM EGTA, 1 mM MgCl_2_, 100 mM KCl, 10% glycerol, 0.05% NP-40, 1 mM DTT, 10 μg/mL leupeptin/pepstatin/chymostatin). Beads were then washed 3 times in wash buffer by rotating for 5 minutes at 4°C. Bound protein was eluted with 100 mM glycine pH 2.6, precipitated by addition of 1/5th volume TCA at 4°C overnight. TCA precipitant was washed 3 times with −20°C acetone and dried using a speed vac.

For quantitative MRPS38 IP-MS experiments involving extended MRPS38, cells were processed as described above, with no detergent included in the lysis buffer to maintain mitochondrial integrity. This approach allowed for the preservation of intact mitochondria, which were then spun out to limit *in vitro* re-association of nucleolar MRPS38 with mitochondrial proteins.

### Mass spectrometry sample preparation

Proteins underwent digestion and purification following a modified S-trap protocol (Protifi, V4.7). The TCA precipitate was dissolved in a lysis buffer composed of 5% SDS and 50 mM TEAB at pH 8.5, then heated to 95°C for 10 minutes with 20 mM DTT to denature the proteins. Subsequently, samples were treated with 40 mM iodoacetamide for 30 minutes at room temperature to alkylate cysteine residues. The mixture was then acidified to a final concentration of 2.5% phosphoric acid. An S-trap binding/wash buffer was added in a volume six times that of the sample, and the solution was loaded onto S-trap mini columns. Centrifugation at 4000×g for 30 seconds was performed, followed by four washes with 150 µL of the binding/wash buffer, each with the same centrifugation conditions. After the final wash, columns were dried by spinning at 4000×g for 1 minute. On-column digestion was carried out overnight at 37°C in a humidified incubator using 20 µL of 50 mM TEAB (pH 8.5) containing 1 µg of trypsin. Peptides were eluted sequentially with 40 µL of 50 mM TEAB containing 0.2% formic acid, followed by a solution of 50% acetonitrile with 0.2% formic acid. The eluted peptides were quantified using the Thermo fluorescent peptide quantification kit, rapidly frozen in liquid nitrogen, and then lyophilized.

### TMT labelling and peptide fractionation

Approximately 1.5 µg of trypsin-digested peptides were dissolved in 50 mM triethylammonium bicarbonate (TEAB) at pH 8.5. Each sample was then labeled with the TMT10plex Isobaric Labeling Reagent Set (Thermo Fisher Scientific, 90111), using a label-to-peptide weight ratio of 30:1, and incubated for 1 hour at room temperature. To quench the labeling reaction, 0.2% hydroxylamine was added, and the mixture was left at room temperature for 15 minutes. The labeled samples were combined on ice, rapidly frozen, and lyophilized. The pooled TMT-labeled peptides were subsequently purified and fractionated using the Pierce High pH Reversed-Phase Peptide Fractionation Kit (Thermo Fisher Scientific, 84868), following the manufacturer’s guidelines for TMT experiments. After fractionation, fractions 1+2, 3+4, 5+6, and 7+8 were pooled, flash-frozen, and lyophilized.

### Mass spectrometry data acquisition

Lyophilized peptides were reconstituted in 0.2% formic acid to achieve a final concentration of 250 ng/µL. Mass spectrometric analysis was conducted using a Thermo Scientific Orbitrap Exploris 480 mass spectrometer, equipped with a FAIMS Pro interface, and coupled to an EASY-nLC chromatography system. Peptide separation was performed on a 25 cm analytical column (PepMap RSLC C18, 3 µm, 100Å, 75 µm inner diameter). The chromatographic gradient was executed at a flow rate of 300 nL/min as follows: 6–21% solvent B over 41 minutes, 21–36% B for 20 minutes, 36– 50% B for 10 minutes, 50–100% B over 15 minutes, followed by column washing and re-equilibration steps. The mass spectrometer and FAIMS interface operated in positive ion mode with a spray voltage of 1800 V and an ion transfer tube temperature set to 270°C. Standard FAIMS resolution settings were applied, utilizing compensation voltages of –50 V and –65 V for the first injection, and –40 V and –60 V for the second. Full scan spectra were acquired in profile mode at a resolution of 120,000, covering a mass-to-charge (m/z) range of 350–1200. The instrument employed an automatic determination of maximum fill time, standard automatic gain control (AGC) target, an intensity threshold of 5×10³, selected charge states of +2 to +5, and a dynamic exclusion duration of 30 seconds.

### Mass spectrometry analysis

Raw data files were processed using Proteome Discoverer version 2.4 (Thermo Fisher Scientific) to identify proteins and peptides. The Sequest HT search engine was employed, using the Homo sapiens protein database (UP000005640) supplemented with EGFP sequences and the MRPS38 N-terminal extension sequence when appropriate. The search parameters allowed for a maximum of two missed trypsin cleavage sites. Mass tolerances were set to 10 ppm for precursor ions and 0.02 Da for fragment ions. The analysis considered several post-translational modifications: dynamic phosphorylation (+79.966 Da on serine, threonine, or tyrosine), dynamic oxidation (+15.995 Da on methionine), dynamic acetylation (+42.011 Da at the N-terminus), dynamic loss of methionine (−131.04 Da at the N-terminal methionine), dynamic loss of methionine with acetylation (−89.03 Da at the N-terminal methionine), and static carbamidomethylation (+57.021 Da on cysteine). For TMT experiments, static modifications included TMT6plex (+229.163 Da at any N-terminus) and TMT6plex (+229.163 Da on lysine residues). Isotope correction factors for TMT 10plex were applied according to the manufacturer’s specifications (Thermo Fisher; product number 90111, lot number VK306786). Peptide identifications were filtered using Percolator to achieve a false discovery rate (FDR) of no more than 0.01. For semi-quantitative IPs, a fixed-value PSM validator was used to filter peptides, retaining only those with a delta Cn ≤ 0.05. Peptides mapping to the MRPS38 N-terminal extension were observed, independent of peptide filtering methods.

### *In vitro* transcription, capping, polyadenylation, and purification

Plasmids containing the reporter gene with a C-terminal GFP tag were amplified via PCR with an oligo with the T7 promoter (5′-TAATACGACTCACTATAGGG-3’) in 5′ oligo to generate templates for *in vitro* transcription. The resulting PCR products were digested with DpnI to remove plasmid template, purified using EconoSpin™ All-in-1 Mini Spin Columns (Epoch Life Science, 1920-250) and eluted with RNase-free water. For transcription, 1 µg of purified PCR product was used in an *in vitro* reaction with the HiScribe T7 High Yield RNA Synthesis Kit (NEB, E2040S) in the presence of 20 U/ml SUPERaseIn. To remove free nucleotides and abortive transcripts, the reaction was passed through P30 columns (Bio-Rad, 732-6251), followed by phenol-chloroform extraction for further purification. The transcribed RNA was then capped using the Vaccinia Capping System (NEB, M2080S) according to the manufacturer’s instructions, purified again using P30 columns, and subjected to phenol-chloroform extraction. To add a poly(A) tail, *Escherichia coli* Poly(A) Polymerase (NEB, M0276) was used, followed by another round of P30 column purification and phenol-chloroform extraction. The final capped and polyadenylated RNA was quantified using a Nanodrop, aliquoted, flash-frozen, and stored at −80 °C.

### mRNA transfections for luciferase assays

Cells were cultured in 24-well plates until reaching approximately 80% confluency before transfection. Transfections were performed using Lipofectamine MessengerMAX (Thermo Fisher, LMRNA008) following the manufacturer’s instructions, with 500 ng of mRNA per well and 1 µL of Lipofectamine MessengerMAX. For luciferase assays, cells were harvested 3 hours after transfection.

### Luciferase assays

Luciferase assays were conducted with the Dual-Luciferase Reporter Assay System (Promega, E1980) following the manufacturer’s instructions. Luminescence was measured using the GloMax Discover Microplate Reader (Promega).

### mRNA transfections for live-cell imaging

Cells were grown to ∼80% confluency in a 12-well plate. Cells were transfected with Lipofectamine MessengerMAX with 1000 ng of mRNA and 2 µl of Lipofectamine MessengerMAX per well. Cells were dislodged from the plate using PBS + 5 mM EDTA and reseeded at 20% density in a 12 well glass bottom plate. 16 hours after reseeding, cells were incubated with 0.1 μg/mL Hoescht for 30 minutes then live imaged as described above.

### 5**′** Rapid Amplification of cDNA Ends (RACE)

RACE reactions were performed with Template Switching RT Enzyme Mix (NEB, M0466) according to manufacturer instructions. 1ug total RNA was used for cDNA preparation for all reactions except for *in vitro* transcribed AKR7A2, where 40ng RNA was used. Reverse-transcription primers were annealed with RNA for 5 minutes at 70°C, followed by reverse transcription and template switching using a template-switching oligo (5′-GCTAATCATTGCAAGCAGTGGTATCAACGCAGAGTACATrGrGrG-3’). For reverse transcription, samples were incubated for 90 minutes at 42°C and 5 minutes at 85°C. PCR amplification was performed with Q5 Hot Start High-Fidelity Master Mix (2X) (NEB, M0494) using 5-20% of cDNA product from reverse-transcription depending on target transcript abundance. To increase specificity, nested primers internal to the reverse-transcription primers were used for PCR amplification, along with a touchdown-PCR protocol with initial cycles of high annealing temperatures (72°C, 70°C) to enrich for target-specific products. PCR-amplified products were run on 1.8% agarose gels for imaging or purified with Zymo DNA Clean and Concentrator kits (D4004) for sequencing analysis using the MGH CCIB DNA core complete amplicon sequencing service. The sequencing reads were assembled de novo, and the number of reads with a unique 5’ end was quantified in the 5’ RACE plots.

### Lentivirus generation

To generate lentivirus, HEK293T cells at ∼80% confluency in a 6 well plate was transfected with 1.2 µg lentiCas9-sgRNA transfer plasmid with puromycin resistance or mCherry, 1 µg psPAX2 packaging plasmid, and 0.4 µg vsFULL envelope plasmid using Xtremegene-9 (Roche). After 16 hours post transfection, the media was swapped. After an additional 24 hours, the lentivirus in the supernatant was collected and stored at - 80°C.

### Competitive growth assays

Fluorescent (eBFP2 or mCherry) HeLa cells were infected with lentivirus carrying Cas9-sgRNA targeting a specific MRPS38 locus (5′-GGCGGCGGCCACAGGTCCCA-3’, 5′-GGGCCCGGGGTCCAGCCGAG-3’, and 5′-AGCGGGAGGCCTGAGGGCGC-3’ for isoform specific guides and 5′-AGGAACGGCCCTCAACAGCT-3’, 5′-GTGTCCGCCCAGCCAGATAG-3’, 5′-GGAAGCTGGTGAAGAAGACG-3’ for all MRPS38 isoforms). Cells at ∼90% confluency were spinfected with lentivirus in the presence of 10 μg/mL polybrene. After 16 hours, the media was swapped. After an additional 24 hours, cells were moved into a new plate and 0.4 μg/mL puromycin was added for 72 hours. After the selection was complete, equal amounts of control eBFP cells were mixed with mCherry cells with Cas9 targeting a specific locus. The population of eBFP:mCherry cells were monitored every 3 days using the BD FACSymphony A1 Cell Analyzer (BD Biosciences).

HeLa cells were infected with lentivirus carrying Cas9-sgRNA targeting specific regions of TRNT1 (Mitochondrial only isoforms: gRNA1: 5′-ACAGGCCAGTGCTGAACCGT-3’ and gRNA2: 5′-GGTGCCTGTATCATTGGCAC-3’; Both isoforms: gRNA1: 5′-GAAGGACTGAAGAGTCTGAC-3’ and gRNA2: 5′-AACTTGCTCTCACCTATATA-3’) or the HS1 locus (5′-GCCGATGGTGAAGTGGTAAG-3’), and mCherry or BFP. Cells at ∼90% confluency in a 6 well plate were spinfected with lentivirus in the presence of 10 μg/mL polybrene. After 16 hours, cells were moved into a 10 cm plate. Cells were grown for another day at 37C, then sorted for mCherry or BFP. After sorting fluorescent cells, the cells were grown for an additional 2 days. Equal amounts of HS1 sgRNA BFP cells were mixed with TRNT1 sgRNA mCherry cells. The population of BFP:mCherry cells were monitored every 3 days using the BD FACSymphony A1 Cell Analyzer (BD Biosciences).

### Total HPG incorporation assay

Polyclonal isoform specific TRNT1 knockout cells were generated as described above. Translation rates were monitored 6 days after lentiviral transduction (or 4 days after sorting). Cells were grown in 12 well plates, incubated with 500 μM Click-IT L-Homopropargylglycine (HPG) for 30 minutes in standard DMEM. Following this incubation, cells were washed once with DMEM, trypsinized, and fixed with 4% formaldehyde for 15 minutes at room temperature. After fixation, cells were washed with PBSTx (PBS + 0.1% Triton X-100) 3 times and blocked with AbDil overnight. Click chemistry was performed by incubating the cells in 0.1M Tris pH 8, 1 mM CuSO4, 0.1M ascorbic acid, and 5 μM Alex Fluor 647 Azide (Thermo, A10277) for 30 minutes in the dark. After click chemistry, cells were washed 3 times with PBSTx, strained into flow cytometry tubes, and analyzed using BD FACSymphony A1 Cell Analyzer (BD Biosciences). HPG signal in these assay measures the summed translation rates of cytosolic and mitochondrial ribosomes.

### Mitochondrial translation assay

Polyclonal isoform specific TRNT1 knockout cells were seeded on poly-Lysine treated coverslips. Mitochondrial translation rates were monitored 6 days after lentiviral transduction (or 4 days after sorting). Cells were in incubated with 500 μM Click-IT L-Homopropargylglycine (HPG) for 30 minutes in standard DMEM. Following this incubation, cells were washed once with DMEM, once with PBS, then fixed with 4% formaldehyde for 15 minutes at room temperature. After fixation, cells were washed with PBSTx (PBS + 0.1% Triton X-100) 3 times and blocked with AbDil overnight. Click chemistry was performed on the coverslip by incubating the cells in 0.1M Tris pH 8, 1 mM CuSO4, 0.1M ascorbic acid, and 5 μM Alex Fluor 647 Azide (Thermo, A10277) for 30 minutes in the dark. After click chemistry, cells were incubated with 0.1 μg/mL Hoescht for 15 minutes, washed 3 times with PBSTx, then mount in PPDM (0.5% p-phenylenediamine and 20 mM Tris-Cl, pH 8.8, in 90% glycerol) and sealed with nail polish. Images were acquired with Deltavision Ultra (Cytiva) system using a 60x/1.42NA objective. Nuclear HPG signal represents product from cytosolic translation whereas mitochondrial signal indicates mitochondrial translation.

### 5**′** UTR length analysis

Human genomic sequences and annotations were downloaded from the GENCODE website (release 25, GRCh38.p7, primary assembly or main annotation). Annotations for protein-coding genes were extracted, and the isoform with the longest ORF was selected to represent that gene in (Fig. 4I; Fig. S4E-F). For analysis in (Fig. S4C, S4G-H) the most highly expressed mRNA isoform in HeLa cells was used in this analysis. The mRNA isoform with the highest transcript per million (TPM) was determined using Kallisto ^71^ using default settings from HeLa cell RNA-sequencing data. GENCODE V25 transcript isoforms were used for Kallisto. MitoCarta3 ^64^ was used to define genes that are mitochondrial.

The transcripts were used for AKR7A2 5′ UTR lengths: human (ENST00000235835), rhesus (ENSMFAT00000078522), mouse (ENSMUST00000073787), and frog (TNeu078h07). We did not use the Ensembl annotation for frog AKR7A2 because the 5′ UTR is repetitive in this annotation.

### GO term enrichment analysis

All GO term analysis was performed using GOrilla ^72^. Two unranked gene lists for this analysis with genes ≤50 nt 5′ UTR and >50 nt 5′ UTR.

### ClinVar analysis and Swissisoform

To facilitate the analysis of alternative protein isoforms and their role in understudied disease variants, we provide SwissIsoform, a computational framework that integrates genomic, transcriptomic, and mutation data with translational isoforms. SwissIsoform takes alternative translation start sites from ribosome profiling experiments and maps them to transcript features, enabling characterization of potential truncated protein isoforms and their relationship with publicly available mutation data.

The SwissIsoform analysis pipeline identifies transcript-truncation pairs where alternative translation initiation sites overlap with annotated transcripts. For each pair, mutations from databases such as ClinVar are analyzed to determine which variants fall within these truncation regions. The framework quantifies mutations by impact type and generates visualizations showing transcript structure, truncation regions, and mutation locations. This approach reveals potentially pathogenic mechanisms associated with alternative translation events, providing insights into how disease-associated variants may disrupt protein function through mechanisms not captured by conventional mutation analysis.

Additionally, to facilitate filtering an extensive list of alternative isoforms for follow-up studies, SwissIsoform generates amino acid sequences for both annotated and alternative isoforms. This capability enables the use of protein language model-based tools, such as DeepLoc, to narrow down variants of interest. SwissIsoform produces standardized outputs that can be directly input into DeepLoc (as done in this manuscript) to identify variants where subcellular localization differs between the annotated protein and the alternative isoform. This integration helps researchers prioritize functionally significant alternative isoforms by highlighting those with potentially altered biological properties, streamlining the selection of candidates for experimental validation.

